# The capacity for adaptation to climate warming in an annual plant (*Brassica rapa*)

**DOI:** 10.1101/2022.10.01.510426

**Authors:** Cameron P. So, Karl Grieshop, Arthur E. Weis

**Author notes:** **Corresponding author:** Cameron P. So; Address: Department of Biology, McGill University, Montreal, QC, Canada.

## Abstract

The persistence of a declining population in the face of environmental change may depend on how fast natural selection restores fitness, a process called “evolutionary rescue”. In turn, evolutionary rescue depends on a population’s adaptive potential. Fisher’s theorem states that a population’s adaptive potential equals the additive genetic variance for fitness (*V_A_*(*W*)) divided by mean fitness 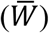. Both the numerator and denominator of this rate can differ across environments even when holding allele frequencies constant. However, little is known about how these rates change in wild populations during adaptation, including changes in additive and dominance variance. We assessed the change in adaptive potential and dominance variance in fitness (*V_D_*(*W*)) for a Québec population of wild mustard (*Brassica rapa*) under climate warming. We also assessed adaptive constraints that could arise from negative genetic correlations across environments. We grew a pedigreed population of 7000 plants under ambient and heated (+4°C) temperatures and estimated the change in 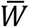, *V_A_*(*W*), *V_D_*(*W*), and the cross-environment genetic correlations (*r_A_*). As predicted, estimates of *V_A_*(*W*) and adaptive potentials were higher under heated conditions but non-significantly so. This is perhaps because, surprisingly, plants exposed to a warmer climate exhibited greater 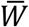. Nevertheless, increased fitness in the warmer environment suggests a plasticity-based short-term potential for adaptation, and that weak but non-significant genetic correlations across environments will enable slow on-going adaptation to warming. Overall, this population of *B. rapa* harbours existing genetic architecture to persist under warmer temperatures through pre-adaptation but not through evolutionary rescue.

## INTRODUCTION

Population persistence can be threatened by environmental change. When unfavourable new conditions reduce the net reproductive rate (*i.e*., absolute mean fitness 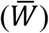) below 1 such that most individuals fail to replace themselves, extinction becomes inevitable. Population decline can be avoided if individuals express phenotypic plasticity in ecologically important traits that maintains fitness. Alternatively, populations could conceivably “evolve their way out” of decline, a process called evolutionary rescue (Bell 2013; Gomulkiewicz and Shaw 2013; Gonzalez and Bell 2013). Rescue in turn, requires sufficient additive genetic variance for fitness (Shaw and Shaw 2014). However, estimates of additive genetic variance for fitness are rare in wild populations (due to logistical and statistical challenges, including expansive pedigrees), and even more seldom estimated in anticipated future environments (Hendry et al. 2018; but see Peschel et al. 2021). Predicting population persistence and evolutionary rescue is therefore limited by the scarcity of estimates of additive genetic variance for fitness across environments.

Rescue prospects of a given population are theoretically predicted from a well understood relationship between phenotype and fitness, and of the genetic architecture underlying phenotypes (Puentes et al. 2016). Yet investigators will seldom, if ever, know all the relevant traits for adaptation to a novel environment. Even if a population harbours high levels of additive genetic variance for the individual traits, the genetic covariances among them alongside other unmeasured trait covariances, make predictions on evolutionary trajectories uncertain (Walsh and Blows 2009). For these reasons, focusing on lifetime fitness directly provides a clearer understanding of on-going adaptation and future population persistence (Shaw and Shaw 2014; Shaw 2019).

If a population harbours at least some individuals with genotypes giving rise to traits and trait combinations suited to the new conditions, then the additive genetic variance in net reproductive rate will be greater than zero. In view of Fisher’s (1930) Fundamental Theorem of Natural Selection, the greater the additive genetic variance in fitness, the faster the rate of adaptation (*i.e*., adaptive potential). The genetically-based change in population mean 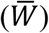 fitness after one generation 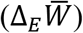 is given as the ratio between additive genetic variance for fitness (*V_A_*(*W*)) and population mean fitness:

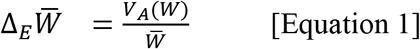

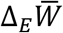 specifically refers to the change in mean fitness caused by evolution, i.e., change in allele frequencies within a generation (Fisher 1930; Ewens 1989; Grafen 2015). The theorem indicates that so long as *V_A_*(*W*) > 0, the mean breeding value for fitness of will increase in the next generation.

Ongoing selection tends to exhaust genetic variation, and so *V_A_*(*W*) is expected to be low (although mutation and gene flow replenish variation). Indeed, the few empirical studies to date indicate *V_A_*(*W*) is typically quite low (Burt 1995; Hendry et al. 2018). As selection pushes favoured alleles toward fixation, additive genetic variance declines. At the same time, recessive alleles may linger, hidden from selection in heterozygotes and thus contributing to *V_D_*(*W*). However, low additive genetic variance in fitness need not imply low genetic variance in the underlying functional traits. Interactions among traits, and their interaction with environment, can mute their genetic contributions to fitness until an environmental change alters these interactions and thereby changes 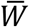 and *V_A_*(*W*), even if underlying allele frequencies are held constant.

Exposing a population to a novel environment can reveal cryptic genetic variance for fitness (Paaby and Rockman 2014; Shaw and Shaw 2014; Zan and Carlborg 2020). This environmental shift could affect *V_A_*(*W*) in two different but non-mutually exclusive ways. First, novel conditions during development can alter the mapping of genotype onto phenotype, changing the mean and/or variance in key traits. The additive effect on phenotype for an allele can increase/decrease with environmental inputs (de Jong 1990). If segregating alleles at a locus differ in their environmental sensitivity, then phenotypes produced by opposite homozygotes will differ more in some environments than in others. Strong stabilizing selection in a constant environment will tend to fix allelic combinations that produce an optimal phenotype in that environment, where cryptic genetic variation remains hidden from selection until/unless revealed by a plastic response to environmental change. Should environmental change alter gene expression, the same allelic combinations could produce an expanded array of phenotypes (Weis and Gorman 1990; Fossen et al. 2018). This exposed hidden variation can have consequent effects on fitness, likely increasing *V_A_*(*W*).

A second way for *V_A_*(*W*) to change with environment is through conversion from dominance and epistatic variance. This conversion occurs when selection goes from stabilizing to directional (Fig. 1). Wright (1934) postulated that the additive genetic variance expressed in a trait near the start of a developmental or metabolic pathway can be converted in part to dominance variance in a downstream fitness-related trait by a non-linear relationship (Gilchrist and Nijhout 2001; Billiard et al. 2021; So et al. in press). Stabilizing selection, by definition, is a non-linear relationship between phenotypic value and fitness. Thus, even if all alleles have additive influence on a key trait, their effect on fitness will show dominance effects (Manna et al. 2011). Figure 1A shows a simple example, where the mean value of trait sits at the optimum. Alleles act additively on the trait but projecting their influence onto fitness via the non-linear fitness function, the alleles show over-dominance for fitness. All variance in fitness is due to genetic differences, yet *V_A_*(*W*) is relatively low. An environmental shift can convert the dominance variance to additive by changing the alignment between the phenotypic mean and optimum. This can occur through a shift in optimal phenotype (Fig. 1B) or through maladaptive phenotypic plasticity in the underlying trait (Fig. 1C). Concordant shifts in the fitness function and the phenotypic mean during a plastic response to environmental change can keep *V_A_*(*W*) low despite promoting local adaptation (Fig. 1D). Hence, the expected change in *V_A_*(*W*) during early adaptation may depend on how closely the plastic response tracks the novel phenotypic optimum.

**Figure 1:**
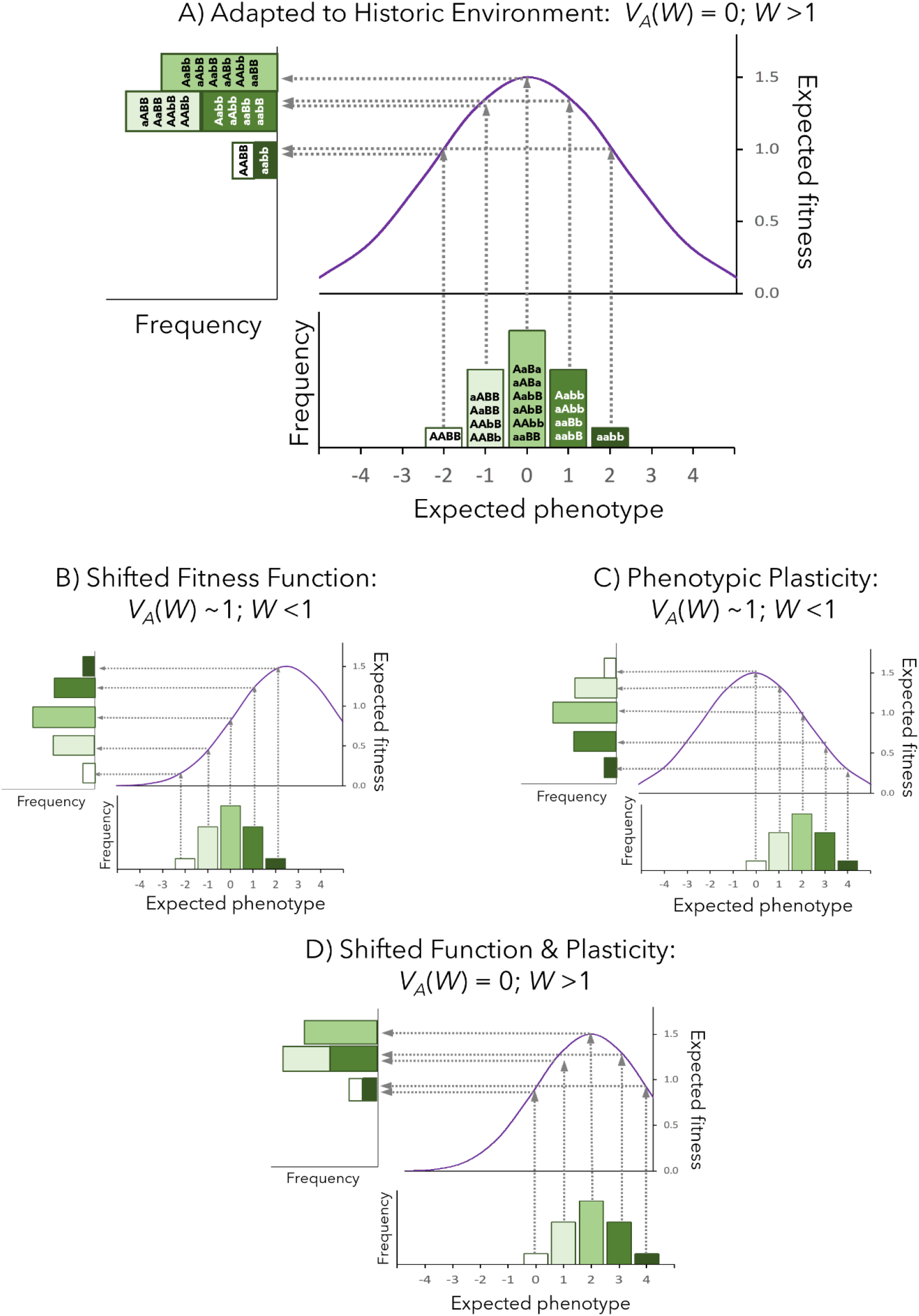
A conceptual illustration of how mean fitness and the additive genetic variance for fitness can change with a shift in the environment. The graph below each section shows the distribution of a phenotype, encoded by different genotypes, corresponding with values of the fitness function above. The integral projection of phenotypic values and its relationship with fitness is a frequency distribution of fitness arranged by genotypic values, shown on the left side. When a population is present in its **(A)** historical environment, *V_A_*(*W*) is low and 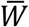 is above 1. However, an **(B)** environmental shift can move the fitness function and rearrange relationships between phenotypes and fitness. This results in a fitness distribution with increased *V_A_*(*W*), lower 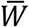, and a greater potential for adaptation. Conversely, **(C)** the developmental environment can induce maladaptive phenotypic plasticity for traits expressed at maturity. For example, a heatwave during the seedling stage of a plant. This increases *V_A_*(*W*) but decreases 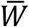 and thus adaptive potential. If an **(D)** an environmental shift changes the fitness function, but the population responds with adaptive plasticity, then *V_A_*(*W*) remains low and 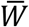 is above 1, similar to historical conditions.

Anthropogenic climate change has propelled wild populations into novel environmental conditions, instigating concern for population persistence under future warming (Parmesan 2006; Hoffmann and Sgrò 2011). In response to multi-year droughts, naturalized Californian populations of *Brassica rapa* (Gulden et al. 2008) have rapidly evolved earlier flowering time and correlated traits (Franks et al. 2007; Franks 2011; Hamann et al. 2018). To evaluate the potential of this annual field mustard to future warming, we estimated the additive genetic variance for fitness (Shaw and Shaw 2014). We used seeds collected in 2008 from Saint-Georges, Québec to produce a pedigreed population from a multi-generational breeding scheme. The F3 experimental generation exhibited varying degrees of known relatedness, from full sibling to half-first cousins, thereby permitting estimation of additive and dominance genetic variation (Lynch and Walsh 1998). We planted the experimental generation into the field at the Koffler Scientific Reserve (KSR) and exposed all family groups to either ambient or heated (+4°C) temperatures.

With our experimental design, we addressed the following questions: (1) Do elevated temperatures decrease the population means of growth performance traits, fitness components and lifetime fitness? (2) Do the additive genetic variances increase under warming. If so, is the increase in *V_A_* through the conversion of dominance genetic variance? If not, is it due to a plastic response that aligns with the novel phenotypic optima? (3) Do genetic correlations among fitness components within and between thermal environments, and in lifetime fitness between environments, impose a constraint on increased adaptation?

## METHODS

### Study system

*Brassica rapa* (field mustard) is an annual, obligately outcrossing, herbaceous plant. Originally introduced from the eastern Mediterranean *via* Europe, the plant has naturalized in locations across North America (Gulden et al. 2008), occupying a broad range of habitats (Warwick et al. 2000) and thereby suggesting high genetic diversity within and among populations (Annisa et al. 2013). Importantly, the self-incompatibility and floral characteristics of *B. rapa* promote estimation of genetic variance components, where desired relationships produced from controlled crosses can increase the power of estimating additive, dominance, and maternal variance.

### Breeding design for experimental population

Using seeds collected in 2008 from a parental population consisting of >5000 plants growing at the margins of a fallow field near Saint-Georges, Québec (46.15N, 70.72W), we conducted 2 generations of controlled greenhouse crosses to produce a 3-generational pedigree (F1 to F3; see **Figs. S1&2** for more information). We generated 62 maternal full-sib families, where individuals of varying degrees of relatedness permitted the estimation of additive (*V_A_*), dominance (*V_D_*), and maternal (*V_M_*) components of genetic variance. Specifically, we partitioned components of variance, including environmental variance (*e*), from the relationship among full-sibs 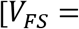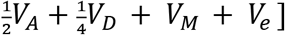 using relationships between reciprocal full siblings [*V*_*R*(*FS*_) = *V_M_* + *V_e_*] among first cousins 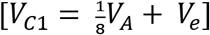, and among half first cousins 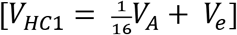 (Lynch and Walsh 1998; Wilson et al. 2010).

### Experimental climate warming array

To evaluate the adaptive potential of *B. rapa* to future climate change, we used the Experimental Climate Warming Array (henceforth referred to as warming arrays) at the Koffler Scientific Reserve (KSR) in King City, Ontario (44.03N, 79.53W). At each experimental plot, six infrared heaters are angled towards the ground to produce a uniform, 3m diameter heat shadow (Kimball et al. 2008; Wadgymar et al. 2015). Using a command-control system, leaf canopy temperatures in both heated and control plots were constantly monitored such that infrared emissions maintained a constant differential of 4°C. Temperatures thus mimicked expected mean global surface temperatures in year 2100 (Intergovernmental Panel on Climate Change 2014). The warming arrays could not be turned on in early spring because sensors were calibrated to only detect leaf temperatures, and so, we installed open top chambers (**Fig. S3**) over all 6 experimental plots for 2 weeks (Intergovernmental Panel on Climate Change 2014). The open top chambers were designed to introduce early spring seasonality, although we recognize that heating the plots directly with infrared prior to emergence may have affected plant development outcomes. Finally, we exposed the experiment to the surrounding biotic environment but excluded large herbivores from the fenced area.

### Experimental planting design

A total of 7270 individuals representing 62 full-sib families (each split into 2 reciprocal maternal sub-groups from reciprocal crosses) were planted at the warming arrays in November 2018 prior to ground freezing. We sowed half of the seeds in the experimental warmed plots (heated) and half in plots without heating (ambient). A week before planting, each plot was covered with a 5 cm layer of seed-free topsoil to prevent stray *B. rapa* seeds from previous experiments germinating. 10 seeds from each of the full-sibships (5 per maternal sub-group) were planted per plot for a total of 605 replicates per plot (N = 5 seeds × ~2 sub-maternal groups × 62 families × 12 plots = 7270 total seeds). Each seed was randomly assigned a position within a 20×30 grid and planted at ~1cm depth with ~2.5 cm spacing between individuals. By the end of the field season, the initial experiment sample size of N=7270 was reduced to N=6809 individuals due to identification issues, accidental damage during handling, and inadequate replication of 5 maternal sub-groups.

### Fitness and trait data collection

We determined measures of lifetime fitness (*W*) from weekly censuses in spring and summer of 2019. The main components of *W* include overwintering survival to flowering (binomial) and fecundity (Poisson distributed). We measured fecundity by counting silique production per plant. Counting the number of seeds per plant, in addition to the male components of fitness, was not practical in an experiment of this size. Several growth performance/functional traits were also measured to determine any correlation to fitness. These include height at flowering, final stem diameter at the root transition zone, the number of leaf nodes, and the number of inflorescences. Height was measured as the length from the root-transition zone to the rachis of the lowest inflorescence.

### Historical weather data

Historical climate data consisting of temperature (°C) and precipitation (mm) was collected from two federally operated weather stations to compare growing season temperature and precipitation at the experimental site (King City, ON) in year 2019 to the parental collection site (Saint-Georges, QC) over the past recorded climate normal (1981-2010). The weather station closest to the field experiment is situated 28 km south of KSR (43.78N, 79.47W), while the station closest to the parental collection site is situated 1.5 km east (46.15N, 70.7W).

### Trait means

We evaluated changes in trait means between environments using R version 3.6.3 (R Core Team 2020) and generalized linear mixed models. Specifically, we used the R package *lme4* version 1.1-21 (Bates et al. 2015) where environment was set as a fixed effect and plot nested under environment as a random effect. For lifetime fitness, we used the R package *glmmTMB* version 1.0.1 (Brooks et al. 2017) to account for zero-inflation. For phenological traits, we used *Cox* mixed effect regression models (Bradburn et al. 2003) from the R package *survival* version 3.1-8 (Therneau and Grambsch 2000). In these models, we tested environment-by-sibship interactions as fixed effects and plot nested under environment as random effects. Only germinated plants were evaluated in flowering phenology assessments.

### Genetic variation

To estimate genetic variance components for each trait of interest, we used the R package *MCMCglmm* version 2.29 (Hadfield 2010), and high-performance computing provided by the *Centre for the Analysis of Genome Evolution* (CAGEF) at the University of Toronto. We fit mixed effect “animal” models with plot set as a fixed effect and additive, dominance, and maternal components as random effects. For estimations of dominance genetic variance, we used the R package *nadiv* version 2.16.2.0 (Wolak 2012) to construct a dominance relatedness matrix. All models were fitted separately for each environment for purposes of extracting environment-specific parameters. After which, we used the R package *QGglmm* version 0.7.4 (de Villemereuil et al. 2016) to transform variance parameters from the latent to data scale for binary and Poisson distributed traits. To calculate mean fitness and thus estimate adaptive potential 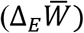, we averaged mean fitness 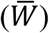 over plot fixed effects (see de Villemereuil et al. 2016, eq. 17). Note that plot was set as a fixed effect to investigate plot-level effects, and later to improve model convergence. For growth and reproductive traits, we signify the rate of adaptation as 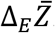. Specifications for each model, which include priors and family distributions used, are provided in the *Supplementary Methods*.

### Genetic correlations

We attempted to estimate cross-environment genetic correlations (*r_A_*), or the degree of shared gene expression between environments because negative *r_A_* indicates selection favouring different trait values between environments, which could disrupt adaptation. Specifically, we estimated the genetic covariance for each trait across environments in a full model (i.e trait ~ environment). These models generally failed model diagnostic tests (e.g poor convergence and high autocorrelation). Hence, we resampled *r_A_* from the stored posterior estimates of F2 parental breeding values from separate within-environment models. This approach addresses the otherwise anticonservative outcomes that can arise when using breeding values in this way by incorporating the uncertainty in those breeding values into the point estimate of *r_A_* and its 95% credibility intervals (Grieshop et al. 2021b). Finally, we also resampled the genetic correlation between survival and fecundity within each environment separately using an approach analogous to that for *r_A_* (above).

## RESULTS

### Experimental climate in relation to historical patterns

The 2019 climatic conditions at the experimental site were similar to the 30-year normal (1981-2010) at the site of the parental source population (Fig. 2). Average daily temperatures throughout the growing season approached historical conditions (Fig. 2A) and precipitation patterns were very similar in the months to which *B. rapa* germinated and underwent growth (Fig. 2B). Altogether, the experimental warming imposed in our study arguably represented the climate scenario of an increased mean temperature by +4°C (Year 2100 RCP8.5, IPCC AR5).

**Figure 2:**
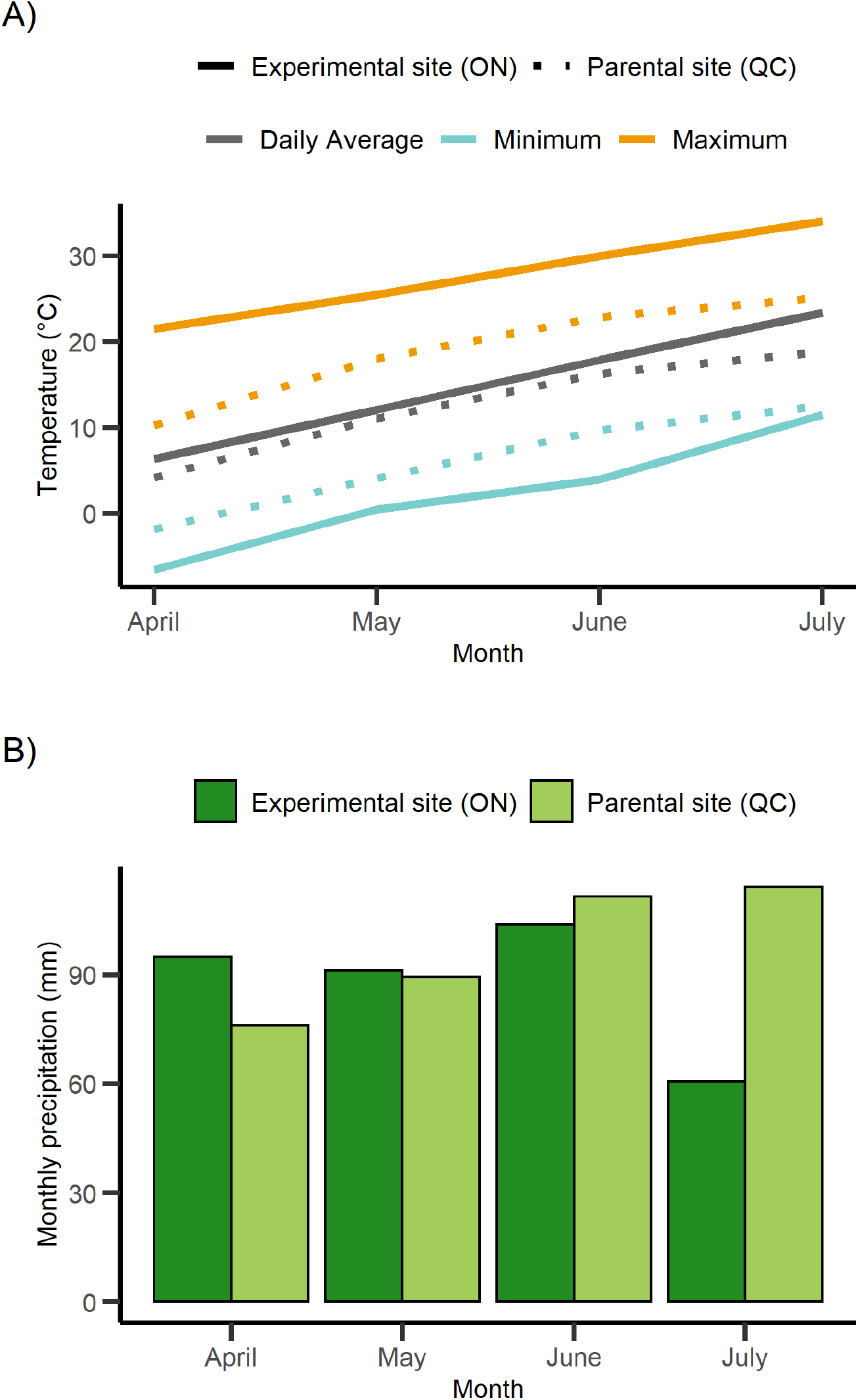
The **(A)** daily temperature and **(B)** monthly precipitation at the experimental site (Koffler Scientific Reserve in Ontario) in the 2019 field season matched the 30-year normal (1981-2010) of the parental site (Saint-Georges in Québec). Thus, the manipulated temperatures in our study should reflect a future climatic regime (+4°C, AR5 IPCC). All climate data retrieved from Environment and Climate Change Canada (ECCC).

### Effects of warming on trait means

All traits expressed greater values on average under the heated environment (**Table 1; Figure 3**). Leaf production and height at flowering was significantly greater under warming while stem diameter was non-significantly greater (**Table 1A**). Warming also initiated earlier germination and flowering time and shifted the flowering season interval (**Fig. S5**). For the fitness components, survival to flowering was similar between environments, however mean fecundity was greater by 48% under warming (**Table 1B**). Lifetime fitness 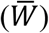 was greater by 61% under heated conditions. Overall, 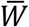 in the heated environment exhibited a greater mean and variance compared to the ambient environment (**Table 1B**).

**Table 1:** A summary of trait means and probabilities per environment, followed with significance testing of the environment using Type II ANOVAs applied for **(A)** growth and reproductive traits and **(B)** lifetime fitness and fitness components. Standard errors are provided following trait means and probabilities (*μ or p*).

| Trait (*i*) | Filter | GLMM family | Ambient $\mu$ or $p$ (SE) | Heated $\mu$ or $p$ (SE) | Type II Wald $\chi^2$ | Environment (P-value) |
| --- | --- | --- | --- | --- | --- | --- |
| **(A) Growth performance traits** |  |  |  |  |  |  |
| Leaf number | Germinated | Gaussian | 6.49<br>(0.19) | 7.81<br>(0.27) | 24.75<sub>(1,3161)</sub> | **6.531e-07** |
| Height (cm) | Germinated | Gaussian | 30.78<br>(2.29) | 37.45<br>(3.24) | 4.24<sub>(1,2873)</sub> | **0.039** |
| Stem diameter (mm) | Germinated | Inverse Gaussian | 1.91<br>(1.08) | 2.19<br>(1.22) | 1.43<sub>(1,2884)</sub> | 0.2308 |
| Inflorescence | Flowered | Poisson | 1.70<br>(0.098) | 2.50<br>(0.14) | 21.21<sub>(1,2902)</sub> | **4.12e-06** |
| **(B) Fitness components** |  |  |  |  |  |  |
| Survival | All data | Bernoulli | 0.41<br>(0.034) | 0.44<br>(0.035) | 0.47<sub>(1,6802)</sub> | 0.49 |
| Fecundity | Flowered | Poisson | 12.04<br>(1.45) | 20.83<br>(2.51) | 6.27<sub>(1,2902)</sub> | **0.012** |
| Lifetime fitness | All data | Zero infl. Neg-bin. | 4.95<br>(0.72) | 9.24<br>(1.35) | 12.17<sub>(1,6802)</sub> | **4.85e-04** |

**Figure 3:**
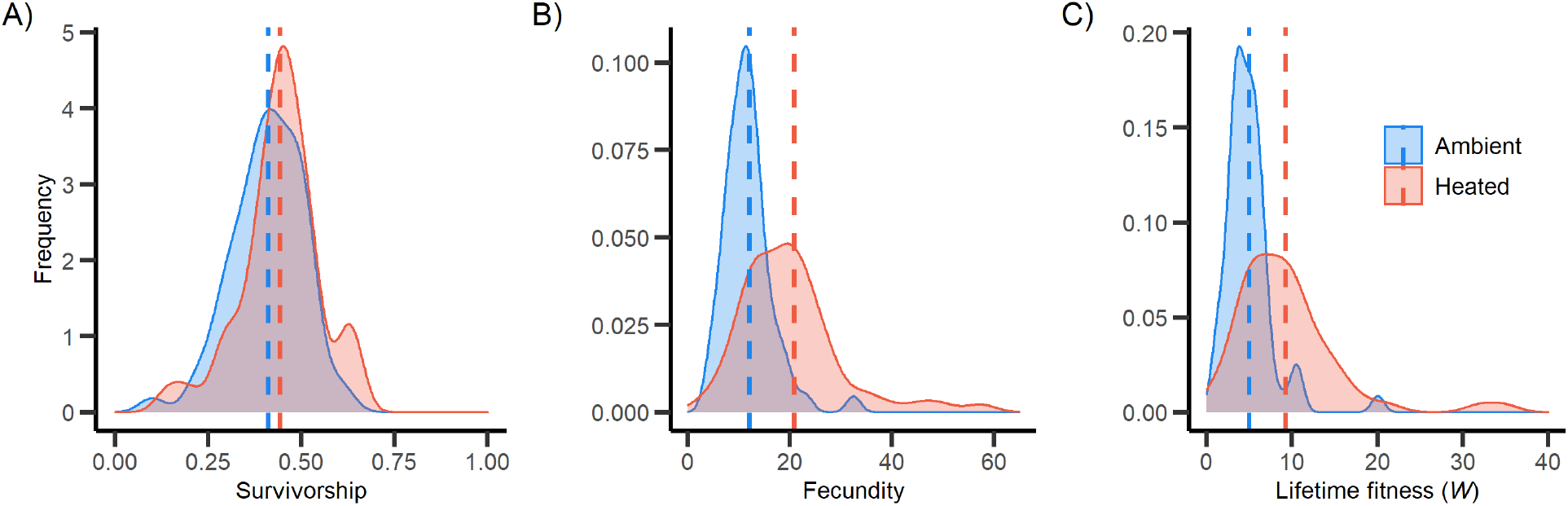
Shifts in phenotypic distributions by heated conditions, calculated using family means, for **(A)** overwintering survivorship to flowering, **(B)** fecundity measured by seed pod number from flowering plants, and **(C)** lifetime fitness measured by total seed pod number. The shift in the distribution of these traits can be compared to the phenotypic frequency distributions outlined in **Figure 1**. Here, the phenotypic distribution shifts low to high variance from the ambient to heated conditions. The mean, represented by the dotted line, also increases under heated conditions for fecundity and lifetime fitness.

### Components of genetic variance

No components of genetic variance significantly differed between the ambient and heated environments for any growth performance traits (**Table 2A**; **Table S2**). Although genetic variance estimates were non-significantly different between environments, we still compare estimates of genetic variance because the *MCMC* sampling process may have biased estimates towards zero (So et al. in press). Additive genetic variance (*V_A_*) for all growth traits were quite low relative to total phenotypic variance (*V_P_*) yet non-significantly greater under heated conditions. In the ambient environment, *V_A_* contributed Dominance genetic variance (*V_D_*) was also greater than *V_A_* across all growth and reproductive traits, although not significantly so due to wide overlapping confidence intervals. Interestingly, *V_D_* contributed substantially to *V_P_* under both ambient (5-20%) and heated conditions (9-16%) while maternal effects were quite negligible compared to other components.

**Table 2:**
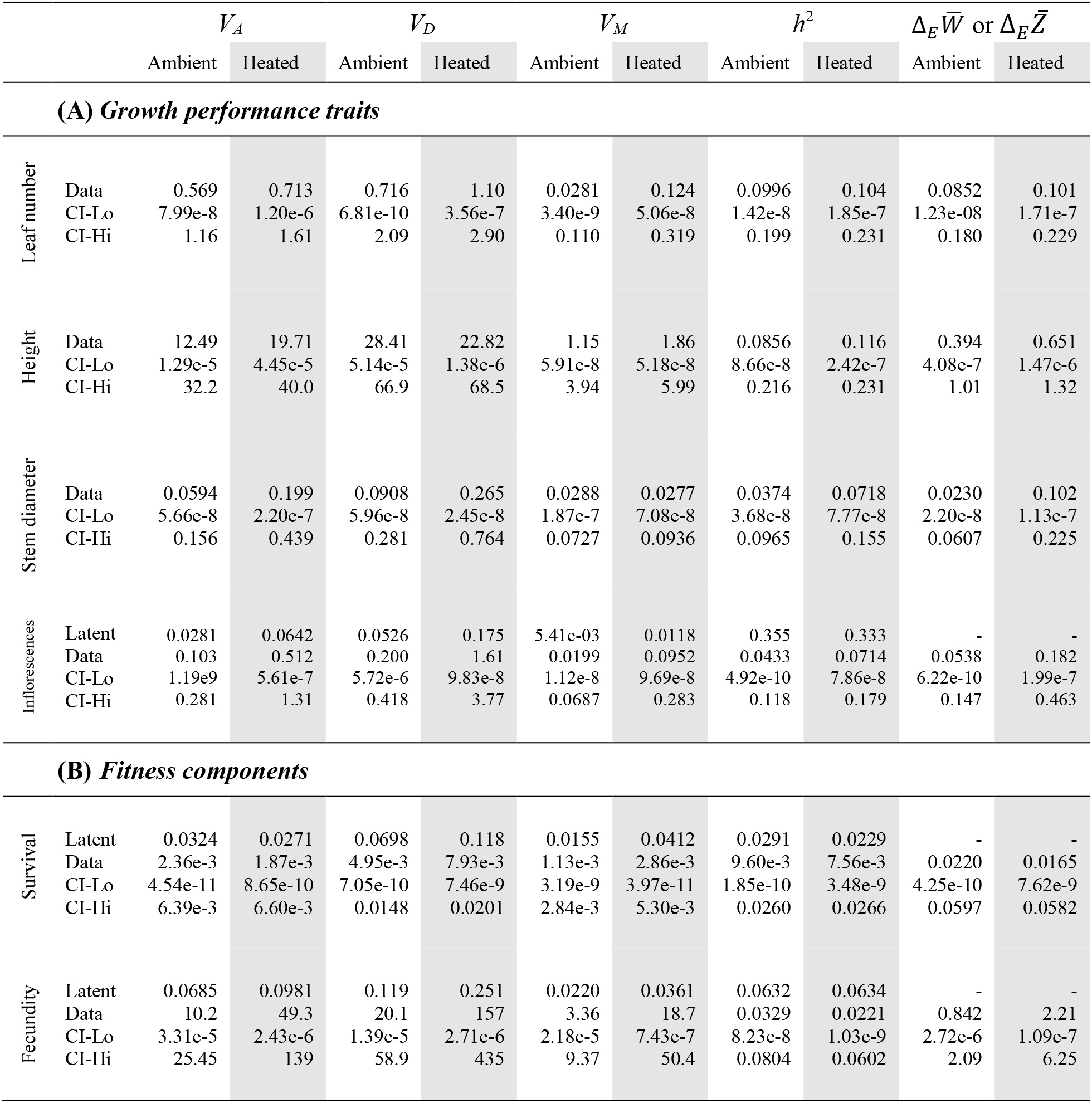
**(A) Growth and reproductive trait**, and **(B) fitness component** estimations of additive genetic variance (*V_A_*), dominance genetic variance (*V_D_*), maternal effects (*V_M_*), narrow-sense heritability (*h^2^*), and adaptive potentials (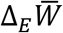 or 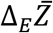. The variances of Bernoulli and Poisson traits are shown on both the latent and data scale, while variances of Gaussian traits are only shown on the data scale. The 95% confidence intervals are only provided on the data scale.

For the fitness components survival and fecundity, *V_A_* did not significantly differ between environments, nor was it different to *V_D_* or *V_M_* within environments (**Table 2B**). We found very low estimates of *V_A_* in both environments on the latent scale, despite substantial “broad-sense” genetic variance measured using sibship means (So et al. in press; **Fig. 3**). When transformed to the data scale, the posterior mean of *V_A_* for survival remained low while fecundity showed a considerable 5-fold increase under heated conditions. Point estimates of *V_D_* were about ~2 times greater than *V_A_* on the latent scale for both survival and fecundity in each environment. When transformed to the data scale, *V_D_* also increased from the latent scale by a factor of 8. Maternal variance was the smallest variance component, and also showed a non-significant increased in heated versus ambient conditions. (Within-family maternal variance is mapped in **Fig. S6**).

Despite posterior mean increases in *V_A_* and *V_D_* under heated conditions, the proportion of additive to phenotypic variance remained similar between environments for survival and fecundity (**Table 2**). Narrow-sense heritability for fecundity only changed from 3.3% to 2.2% between the ambient and heated condition. The proportion of dominance variance to phenotypic variance also remained similar between environments. Ambient *V_D_* accounted for 2% and 6% of the phenotypic variance for survival and fecundity, respectively, compared to 3% and 6.5% in the heated environment. These results indicate no conversion of dominance to additive genetic variance, and that the (non-significant) changes in genetic variance estimates are due to scaling of the mean.

The adaptive potential, 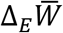 [Eqn. 1], for the fitness components did not significantly differ between environments. Looking solely at posterior means, 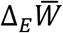 for fecundity was greater by ~2-fold on the data scale in the heated environment 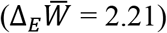 compared to ambient 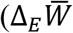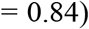. Nevertheless, we found no difference in the potential for adaptation for fecundity under the heated environment. We did not estimate components of genetic variance or adaptive potentials for lifetime fitness because of its zero-inflated Poisson distribution. This statistical challenge can be overcome using *aster* models for estimating *V_A_*(*W*) (Geyer et al. 2007, 2013), however the *aster* approach is currently infeasible for estimating multiple components of genetic variance.

### Genetic correlations between environments

Genetic correlations for traits expressed across ambient and heated environments (*r_A_*) were low and non-significant (**Table 3**; **Table S3**). Confidence intervals overlapped zero with both positive and negative bounds. We were unable to estimate the cross-environment *r_A_* for lifetime fitness because of its zero-inflated Poisson distribution. Reaction norms of sibship means for survival and fecundity indicated limited re-ranking of genotypes and an average slope near zero (**Fig. S7**). While some sibships had positive slopes, the overall reaction norm agreed with low and non-significant estimates of *r_A_*. Interestingly, these non-significant cross-environment genetic correlations contrast significant effects of environment on phenotypic values (*i.e*., contrast strong phenotypic correlations). We also found weak and non-significant genetic correlations between survival and fecundity under both environments (**Table S4**). Note that the bivariate model to estimate the genetic correlation between fitness components included *all* plants in the study, contrasting models of fecundity which included only plants that survived to reproduction. Fecundity in this case is evaluated for each seed sown into the ground and thus differs from the definitions used previously in the univariate model.

**Table 3:** Cross-environment additive genetic correlations and *p* values signifying any differences of *r* from 0 for **(A)** growth and reproductive traits, and **(B)** survival and reproductive components. Each correlation coefficient is provided with upper and lower 95% confidence intervals in parentheses.

| Trait | Cross-environment<br>genetic correlation ($r_A$) | Significant $r$<br>(P-value) |
| --- | --- | --- |
| **(A) Growth performance traits** |  |  |
| Leaf number | 0.129<br>(-0.0925, 0.324) | 0.315 |
| Height | 0.0932,<br>(-0.101, 0.335) | 0.349 |
| Stem diameter | 0.112,<br>(-0.134, 0.274) | 0.497 |
| Inflorescences | 0.0454<br>(-0.136, 0.269) | 0.510 |
| **(B) Fitness components** |  |  |
| Survival | 0.00158,<br>(-0.158, 0.218) | 0.792 |
| Fecundity | 0.0931,<br>(-0.142, 0.275) | 0.525 |

## DISCUSSION

The persistence of a population experiencing environmental change depends upon the population’s adaptive potential, or the rate of change for fitness described as 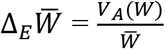 [Eqn. 1] (Fisher 1930; Frank and Slatkin 1992; Shaw and Shaw 2014). Despite the urgency of predicting population persistence to anthropogenic climate warming, estimates of additive genetic variance for fitness (*V_A_*(*W*)) in ratio to mean fitness 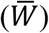 are rare. We exposed a pedigreed population of *Brassica rapa* to ambient and heated (+4°C) temperatures to estimate the change in mean fitness 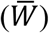, additive genetic variance for fitness (*V_A_*(*W*)), and adaptive potential 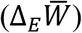 to future warming. We also evaluated whether changes in *V_A_* was due to a conversion of dominance, and whether genetic correlations could constrain the adaptive process. While we decomposed components of genetic variance for each trait as well as survival and fecundity (**Table 2**), we were not able to analyze lifetime fitness because of statistical limitations (see Results). Expansive confidence intervals for genetic variance parameters yielded no significant differences between environments. We attribute this result to either: a pedigree structure lacking replication in first and double cousin relationships, the *MCMC* sampling process pulling estimates towards zero, or truly low estimates of genetic variation (So et al. in press). We interpret our results in review of the three questions highlighted earlier in the sections below.

### Fitness and traits respond plastically to warming

Plants responded positively to warming by increasing trait means and ultimately, lifetime fitness (**Table 1**; **Fig. 3**). Advanced germination time, likely from earlier snowmelt (Henry et al. 2015), translated into earlier resource acquisition, greater growth, and thus improved reproduction. Similar plant responses (Mohan et al. 2019) were found in both a warming experiment of *Arabidopsis thaliana* (Springate and Kover 2014) and a long-term observational study of *Boechera stricta* over a period of climate change-induced warming (Anderson et al. 2012; Mohan et al. 2019). In both studies, warming selected for early flowering which increased fitness. With respect to *Brassica rapa*, evolutionary responses to warming have already been documented with early flowering conferring higher fitness (Franks et al. 2007; Hamann et al. 2018). Although rapid evolution may be adaptive, plasticity in traits such as germination and flowering time can also buffer populations from detrimental environmental effects by maintaining, or even increasing fitness (Matesanz et al. 2010; Chevin et al. 2013; Kopp and Matuszewski 2014; Chevin and Hoffmann 2017; Fox et al. 2019). Lifetime fitness 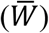 plastically increased in response to warming in our study, but this means the potential for adaptation can also decrease through Fisher’s Fundamental Theorem of Natural Selection 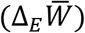. Whether this adaptive potential reduces under warming depends upon the magnitude of additive genetic variance for fitness.

### Additive genetic variance and adaptive potentials

For all traits, we estimated low additive genetic variance (*V_A_*) that did not significantly differ between environments (**Table 2**). Although, it should be noted that we did find (statistically non-significant, but noticeably larger) increases in *V_A_* under the warmer environment when we transformed non-Gaussian parameters (*e.g*., for fecundity and inflorescences) to the data scale (de Villemereuil et al. 2016; Bonnet et al. 2019). Increases in *V_A_* under non-optimal conditions is consistent with other plant studies in the wild (Etterson 2004; Winn 2004; Sheth et al. 2018; Kulbaba et al. 2019; Torres-Martínez et al. 2019) Nevertheless, additive genetic variances translated into low heritability and rates of adaptation for all traits, suggesting that while *B. rapa* may be able to respond plastically, it has a low potential to rapidly evolve to climate warming.

From the limited number of reported *V_A_* estimates in wild populations, our low *V_A_* estimates appear to be uncommon however not without precedent (Hendry et al. 2018). For example, McFarlane et al. (2014) found low *V_A_* for male and female fitness components and for lifetime reproductive success in North American red squirrel populations. In our study, a possible explanation for low *V_A_* estimates could be a population bottleneck in conjunction to founder effects (Whitlock et al. 2002; Castillo et al. 2018). Naturalized populations of *Brassica rapa* found in North America were originally introduced from Europe and have historical roots stemming from the Mediterranean and Central Asia (Ignatov et al. 2010; Guo et al. 2014). The population sampled from Saint-Georges, Québec may have descended from a series of genetic bottlenecks, starting with the introductions from Europe, and subsequently in newly disturbed sites such as roadsides, croplands, and weedy fields. In addition, weed control measures at the Québec site may have at some point reduced population size, causing a loss of allelic variation. Comparing the genetic variation of this widely distributed species across its geographical range would resolve this hypothesis, as plant introductions across continents and regions do not always constitute depressions in genetic diversity underpinning adaptive potentials (McGoey and Stinchcombe 2021).

Even if *B. rapa* was introduced hundreds of years ago allowing opportunity for genetic variation to change, species distribution patterns (e.g season length) could also explain low additive genetic variation (Sheth and Angert 2016; Pironon et al. 2017). The population found in Saint-Georges is likely to be at the leading edge of *B. rapa*’s geographical distribution, stopping just short of the Boreal Forest located northward. Strong directional selection on flowering time at northern range edges (Munguía-Rosas et al. 2011) in addition to directional selection acting on other traits related to fitness can deplete the genetic variation for these traits. Thus, considering the historical and geographical context of plant populations is crucial for understanding why standing additive genetic variance for fitness-related traits can be as low as observed.

### Expression of dominance genetic variance

When populations are exposed to novel environments, the proportional effect of dominance genetic variance (*V_D_*) contributing to fitness variance is expected to diminish (Fig. 1; Cockerham and Tachida 1988; Willis and Orr 1993). In our study, *V_D_* for the fitness components did not significantly change between environments (**Table 2**). In other words, dominance did not convert into additive variance. Surprisingly however, *V_D_* was consistently similar to *V_A_* in magnitude across all traits and was generally ~1.5 – 2-fold greater (and ~5-fold greater for fecundity). These findings are consistent with a recent review of dominance variance in wild populations, in which *V_A_* and *V_D_* of life history traits contributed equally to the total phenotypic variance (Wolak and Keller 2014). The persistence and strength of *V_D_* relative to *V_A_* suggests dominance plays an important role in adaptation (Crnokrak and Roff 1995; Wolak and Keller 2014). For instance, non-additive genetic effects contribute substantially to the fitness component variation in a marine tubeworm (Chirgwin et al. 2017) and excluding dominance would overestimate adaptive potentials (Wilson 2008; Class and Brommer 2020; Clo and Opedal 2021). Further, dominance variance could reflect dominance reversals (e.g., between sexes; Grieshop and Arnqvist 2018), which have been putatively demonstrated between environments (Posavi et al. 2014), and which could potentially mediate plasticity (reviewed in Grieshop et al. 2021a).

### Genetic correlations within and among environments

Life history theory predicts that negative genetic correlations between survival and fecundity can impede the rate of adaptation (Stearns 1989, 1992). In face of climate warming, the magnitude and direction of genetic correlations between life history traits may also change and thus affect rates of adaptation (Sgrò and Hoffmann 2004; Chevin 2013; Wood and Brodie 2015). It is likely that climate change will select for genetical lines that confer high survival and high fecundity. We observed low genetic correlations between survival and fecundity under both environments. This suggests that strong selection for greater survival will have low to no effect on fecundity, and vice versa. Under climate warming, both fitness components could evolve without constraint, and if the true cross-environment correlations are positive it could facilitate rather than constrain adaptation. The cumulative weakly positive cross-environment genetic correlations observed among all traits and fitness components in our study could facilitate a multivariate adaptive response to climate warming, but we were unable to estimate the full multivariate variance-covariance matrix with our data. We note that the synergistic effect of multiple stressors caused by climate change (*e.g*. drought, pollinator abundance) could also alter other traits underpinning fitness components, thereby altering these genetic correlations, including exacerbating and/or alleviating constraints. Researchers should explore how genetic correlations between functional traits change with environmental shifts induced by climate change (Etterson and Shaw 2001; Berger et al. 2014; Puentes et al. 2016).

Genetic correlations within traits and across environments indicate the extent of phenotypic plasticity. We found low cross-environment genetic correlations for all traits (**Table 3**), suggesting differential gene expression under each environment. Importantly, low genetic correlations, which are explained by low additive genetic covariances (*COV_A_*), indicate low adaptive potential for plasticity (Schlichting 1986; Via and Lande 1987; Acasuso-Rivero et al. 2019). Genetic variance for plasticity can be inherently low after adaptive responses to variable environments (*e.g*., pre-adaptation from historical exposure). Alternatively, the positive effect of warming may not have revealed substantial genetic variation to detect genetic correlations. By design, our study was unable to elucidate which explanation is possible. Exposing plants to multiple environments in future studies may provide a better answer (e.g Ensing & Eckert, 2019; Salinas et al., 2019; Torres-Martínez et al., 2019). Understanding how plasticity will adapt in response to increasing temperatures and variability will be central to predicting plant evolutionary responses to contemporary climate change (Jentsch et al. 2007; Franks et al. 2014).

### Conclusion and Future Directions

Our study adds to growing assessments of adaptive potentials in wild populations, through estimations of additive genetic variation, to environmental change (e.g Chapuis et al. 2021; Peschel et al. 2021). We found low additive genetic variance (*V_A_*) for functional traits and fitness components under both ambient and heated environments. Low *V_A_* translated into low adaptive potentials, either represented by heritability or the Fisher’s Fundamental Theorem of Natural Selection (FTNS). Nevertheless, a low potential to adapt does not necessarily equal poor prospects for population persistence. Under heated temperatures, plants grew to greater heights, produced more leaves, and yielded a greater number of seed pods. These findings indicate that *B. rapa* is pre-adapted to climate warming and that phenotypic plasticity will be instrumental to its adaptive capacity. Further work is needed to estimate the *V_A_* for lifetime fitness, especially where dominance is partitioned from genetic variance. Estimates of *V_A_* for lifetime fitness should provide more complete knowledge on the prospects for evolutionary rescue, especially as extreme climate events are expected to severely reduce population growth rate (Hall et al., *in prep*). Still, our work provides immediate implications on the potential for populations to adapt to future climate warming (see Fig. 1D; Anderson & Song, 2020; Shaw & Etterson, 2012), as populations may be pre-adapted but unable to adaptively evolve to warmer conditions.

## ACKNOWLEDGEMENTS

We thank the many volunteers who assisted this study. We are especially grateful for members of the Weis Lab. Notably, S. Rotman attempted the first trial of this experiment and M. Sibolibane collected a large portion of the data. We thank M. Wolak and P. de Villemereuil for their statistical advice on *MCMCglmm*, and P.L. Maurizio for providing his code to run *MCMCglmm* models in parallel. This work was supported with funding from the Koffler Scientific Reserve at Joker’s Hill (CPS, AEW), the Centre for Global Change Science (CPS), the Natural Sciences and Engineering Research Council of Canada (RGPIN-2016-06540 to AEW), and the Swedish Research Council (2018-06775 to KG). The authors declare no conflicts of interest.

## AUTHOR CONTRIBUTIONS

AEW conceived the study, CPS and AW designed the study, CPS performed the experiment, conducted the statistical analyses, and drafted the initial version of the manuscript. KHG provided code and advice on the statistical analyses. CPS and AEW produced the figures. All authors contributed to editing the manuscript.

## DATA AVAILABILITY STATEMENT

R scripts and data used will be made available on the *Dryad Digital Repository*.

## SUPPLEMENTARY INFORMATION

Additional supporting information may be found online in the Supporting Information document.

## REFERENCES

Acasuso-Rivero, C., C. J. Murren, C. D. Schlichting, and U. K. Steiner. 2019. Adaptive phenotypic plasticity for life-history and less fitness-related traits. Proceedings of the Royal Society B: Biological Sciences 286:20190653.

Anderson, J. T., D. W. Inouye, A. M. McKinney, R. I. Colautti, and T. Mitchell-Olds. 2012. Phenotypic plasticity and adaptive evolution contribute to advancing flowering phenology in response to climate change. Proceedings of the Royal Society B: Biological Sciences 279:3843–3852.

Anderson, J. T., and B. Song. 2020. Plant adaptation to climate change—Where are we? J Syst Evol 58:533–545.

Annisa, S. Chen, and W. A. Cowling. 2013. Global genetic diversity in oilseed Brassica rapa. Crop Pasture Sci 64:993–1007.

Bates, D., M. Mächler, B. Bolker, and S. Walker. 2015. Fitting linear mixed-effects models using lme4. J Stat Softw 67.

Bell, G. 2013. Evolutionary rescue and the limits of adaptation. Philosophical Transactions of the Royal Society B: Biological Sciences 368:20120080.

Berger, D., K. Grieshop, M. I. Lind, J. Goenaga, A. A. Maklakov, and G. Arnqvist. 2014. Intralocus sexual conflict and environmental stress. Evolution (N Y) 68:2184–2196. John Wiley & Sons, Ltd.

Billiard, S., V. Castric, and V. Llaurens. 2021. The integrative biology of genetic dominance. Biological Reviews 96:2925–2942. John Wiley & Sons, Ltd.

Bonnet, T., M. B. Morrissey, and L. E. B. Kruuk. 2019. Estimation of genetic variance in fitness, and inference of adaptation, when fitness follows a log-normal distribution. Journal of Heredity 110:383–395.

Bradburn, M. J., T. G. Clark, S. B. Love, and D. G. Altman. 2003. Survival analysis part II: Multivariate data analysis – an introduction to concepts and methods. Br J Cancer 89:431–436.

Brooks, M. E., K. Kristensen, K. J. van Benthem, A. Magnusson, C. W. Berg, A. Nielsen, H. J. Skaug, M. Mächler, and B. M. Bolker. 2017. glmmTMB Balances Speed and Flexibility Among Packages for Zero-inflated Generalized Linear Mixed Modeling. R J 9:378.

Burt, A. 1995. The evolution of fitness. Evolution (N Y) 49:1–8.

Castillo, J. M., B. Gallego-Tévar, E. Figueroa, B. J. Grewell, D. Vallet, H. Rousseau, J. Keller, O. Lima, . Dréano, A. Salmon, and M. Aïnouche. 2018. Low genetic diversity contrasts with high phenotypic variability in heptaploid Spartina densiflora populations invading the Pacific coast of North America. Ecol Evol 8:4992–5007.

Chapuis, M. P., B. Pélissié, C. Piou, F. Chardonnet, C. Pagès, A. Foucart, E. Chapuis, and H. Jourdan-Pineau. 2021. Additive genetic variance for traits least related to fitness increases with environmental stress in the desert locust, Schistocerca gregaria. Ecol Evol 11:13930–13947. John Wiley & Sons, Ltd.

Chevin, L.-M. 2013. Genetic constraints on adaptation to a changing environment. Evolution (N Y) 67:708–721.

Chevin, L.-M., R. Gallet, R. Gomulkiewicz, R. D. Holt, and S. Fellous. 2013. Phenotypic plasticity in evolutionary rescue experiments. Philosophical Transactions of the Royal Society B: Biological Sciences 368:20120089.

Chevin, L.-M., and A. A. Hoffmann. 2017. Evolution of phenotypic plasticity in extreme environments. Philosophical Transactions of the Royal Society B: Biological Sciences 372:20160138.

Chirgwin, E., D. J. Marshall, C. M. Sgrò, and K. Monro. 2017. The other 96%: Can neglected sources of fitness variation offer new insights into adaptation to global change? Evol Appl 10:267–275.

Class, B., and J. E. Brommer. 2020. Can dominance genetic variance be ignored in evolutionary quantitative genetic analyses of wild populations? Evolution (N Y) 74:1540–1550. John Wiley & Sons, Ltd.

Clo, J., and Ø. H. Opedal. 2021. Genetics of quantitative traits with dominance under stabilizing and directional selection in partially selfing species. Evolution (N Y) 75:1920–1935. Evolution.

Cockerham, C. C., and H. Tachida. 1988. Permanency of response to selection for quantitative characters in finite populations. Proceedings of the National Academy of Sciences 85:1563–1565.

Crnokrak, P., and D. A. Roff. 1995. Dominance variance: associations with selection and fitness. Heredity (Edinb) 75:530–540.

de Jong, G. 1990. Genotype-by-environment interaction and the genetic covariance between environments: multilocus genetics. Genetica 81:171–177.

de Villemereuil, P., H. Schielzeth, S. Nakagawa, and M. B. Morrissey. 2016. General methods for evolutionary quantitative genetic inference from generalized mixed models. Genetics 204:1281–1294.

Ensing, D. J., and C. G. Eckert. 2019. Interannual variation in season length is linked to strong co-gradient plasticity of phenology in a montane annual plant. New Phytologist 224:1184–1200.

Etterson, J. R. 2004. Evolutionary potential of Chamaecrista fasciculata in relation to climate change. II. genetic architecture of three populations reciprocally planted along an environmental gradient in the great plains. Evolution (N Y) 58:1459–1471.

Etterson, J. R., and R. G. Shaw. 2001. Constraint to adaptive evolution in response to global warming. Science (1979) 294:151–154.

Ewens, W. J. 1989. An interpretation and proof of the fundamental theorem of natural selection. Theor Popul Biol 36:167–180.

Fisher, R. A. 1930. The genetical theory of natural selection. Clarendon Press, Oxford.

Fossen, E. I. F., C. Pélabon, and S. Einum. 2018. An empirical test for a zone of canalization in thermal reaction norms. J Evol Biol 31:936–943. John Wiley & Sons, Ltd.

Fox, R. J., J. M. Donelson, C. Schunter, T. Ravasi, and J. D. Gaitán-Espitia. 2019. Beyond buying time: the role of plasticity in phenotypic adaptation to rapid environmental change. Philosophical Transactions of the Royal Society B: Biological Sciences 374:20180174.

Frank, S. A., and M. Slatkin. 1992. Fisher’s fundamental theorem of natural selection. Trends Ecol Evol 7:92–95.

Franks, S. J. 2011. Plasticity and evolution in drought avoidance and escape in the annual plant Brassica rapa. New Phytologist 190:249–257.

Franks, S. J., S. Sim, and A. E. Weis. 2007. Rapid evolution of flowering time by an annual plant in response to a climate fluctuation. Proceedings of the National Academy of Sciences 104:1278–1282.

Franks, S. J., J. J. Weber, and S. N. Aitken. 2014. Evolutionary and plastic responses to climate change in terrestrial plant populations. Evol Appl 7:123–139.

Geyer, C. J., C. E. Ridley, R. G. Latta, J. R. Etterson, and R. G. Shaw. 2013. Local adaptation and genetic effects on fitness: Calculations for exponential family models with random effects. Ann Appl Stat 7:1778–1795.

Geyer, C. J., S. Wagenius, and R. G. Shaw. 2007. Aster models for life history analysis. Biometrika 94:415–426.

Gilchrist, M. A., and H. F. Nijhout. 2001. Nonlinear developmental processes as sources of dominance. Genetics 159:423–432. Oxford Academic.

Gomulkiewicz, R., and R. G. Shaw. 2013. Evolutionary rescue beyond the models. Philosophical Transactions of the Royal Society B: Biological Sciences 368:20120093.

Gonzalez, A., and G. Bell. 2013. Evolutionary rescue and adaptation to abrupt environmental change depends upon the history of stress. Philosophical Transactions of the Royal Society B: Biological Sciences 368:20120079.

Grafen, A. 2015. Biological fitness and the fundamental theorem of natural selection. American Naturalist, doi: 10.1086/681585.

Grieshop, K., and G. Arnqvist. 2018. Sex-specific dominance reversal of genetic variation for fitness. PLoS Biol 16:1–21.

Grieshop, K., E. K. H. Ho, and K. R. Kasimatis. 2021a. Dominance reversals, antagonistic pleiotropy, and the maintenance of genetic variation. arXiv preprint arXiv:2109.01571.

Grieshop, K., P. L. Maurizio, G. Arnqvist, and D. Berger. 2021b. Selection in males purges the mutation load on female fitness. Evol Lett 5:328–343. John Wiley & Sons, Ltd.

Gulden, R. H., S. I. Warwick, and A. G. Thomas. 2008. The biology of Canadian weeds. 137. Brassica napus L. and B. rapa L. Canadian Journal of Plant Science 88:951–996.

Guo, Y., S. Chen, Z. Li, and W. A. Cowling. 2014. Center of origin and centers of diversity in an ancient crop, Brassica rapa (Turnip Rape). Journal of Heredity 105:555–565.

Hadfield, J. D. 2010. MCMC methods for multi-response generalized linear mixed models: The MCMCglmm R package. J Stat Softw 33:1–22.

Hamann, E., A. E. Weis, and S. J. Franks. 2018. Two decades of evolutionary changes in Brassica rapa in response to fluctuations in precipitation and severe drought. Evolution (N Y) 72:2682–2696.

Hendry, A. P., D. J. Schoen, M. E. Wolak, and J. M. Reid. 2018. The contemporary evolution of fitness. Annu Rev Ecol Evol Syst 49:457–476.

Henry, H. A. L., J. S. Hutchison, M. K. Kim, and B. D. McWhirter. 2015. Context matters for warming: Interannual variation in grass biomass responses to 7 years of warming and N addition. Ecosystems 18:103–114.

Hoffmann, A. A., and C. M. Sgrò. 2011. Climate change and evolutionary adaptation. Nature 470:479–485.

Ignatov, A. N., A. M. Artemyeva, and K. Hida. 2010. Origin and expansion of cultivated Brassica rapa in Eurasia: Linguistic facts. P. *in* Acta Horticulturae.

Intergovernmental Panel on Climate Change. 2014. Climate Change 2013 - The Physical Science Basis. Cambridge University Press, Cambridge.

Jentsch, A., J. Kreyling, and C. Beierkuhnlein. 2007. A new generation of climate-change experiments: events, not trends. Front Ecol Environ 5:365–374.

Kimball, B. A., M. M. Conley, S. Wang, X. Lin, C. Luo, J. Morgan, and D. Smith. 2008. Infrared heater arrays for warming ecosystem field plots. Glob Chang Biol 14:309–320.

Kopp, M., and S. Matuszewski. 2014. Rapid evolution of quantitative traits: Theoretical perspectives. Evol Appl 7:169–191.

Kulbaba, M. W., S. N. Sheth, R. E. Pain, V. M. Eckhart, and R. G. Shaw. 2019. Additive genetic variance for lifetime fitness and the capacity for adaptation in an annual plant. Evolution (N Y) 73:1746–1758.

Lynch, M., and B. Walsh. 1998. Genetics and analysis of quantitative traits. Sinauer Associates, Inc., Sunderland, MA, MA.

Manna, F., G. Martin, and T. Lenormand. 2011. Fitness landscapes: An Alternative theory for the dominance of mutation. Genetics 189:923–937. Oxford Academic.

Matesanz, S., E. Gianoli, and F. Valladares. 2010. Global change and the evolution of phenotypic plasticity in plants. Ann N Y Acad Sci 1206:35–55.

McFarlane, S. E., J. C. Gorrell, D. W. Coltman, M. M. Humphries, S. Boutin, and A. G. McAdam. 2014. Very low levels of direct additive genetic variance in fitness and fitness components in a red squirrel population. Ecol Evol 4:1729–1738.

McGoey, B. v., and J. R. Stinchcombe. 2021. Introduced populations of ragweed show as much evolutionary potential as native populations. Evol Appl 14:1436–1449.

Mohan, J. E., S. M. Wadgymar, D. E. Winkler, J. T. Anderson, P. T. Frankson, R. Hannifin, K. Benavides, L. M. Kueppers, and J. M. Melillo. 2019. Plant reproductive fitness and phenology responses to climate warming: Results from native populations, communities, and ecosystems. Pp. 61–102 *in* Ecosystem Consequences of Soil Warming. Elsevier.

Munguía-Rosas, M. A., J. Ollerton, V. Parra-Tabla, and J. A. De-Nova. 2011. Meta-analysis of phenotypic selection on flowering phenology suggests that early flowering plants are favoured. Ecol Lett 14:511–521.

Paaby, A. B., and M. v. Rockman. 2014. Cryptic genetic variation: evolution’s hidden substrate. Nat Rev Genet 15:247–258.

Parmesan, C. 2006. Ecological and evolutionary responses to recent climate change. Annu Rev Ecol Evol Syst 37:637–669.

Peschel, A. R., E. L. Boehm, and R. G. Shaw. 2021. Estimating the capacity of Chamaecrista fasciculata for adaptation to change in precipitation. Evolution (N Y) 75:73–85.

Pironon, S., G. Papuga, J. Villellas, A. L. Angert, M. B. García, and J. D. Thompson. 2017. Geographic variation in genetic and demographic performance: new insights from an old biogeographical paradigm. Biological Reviews 92:1877–1909.

Posavi, M., G. W. Gelembiuk, B. Larget, and C. E. Lee. 2014. Testing for beneficial reversal of dominance during salinity shifts in the invasive copepod Eurytemora affinis, and implications for the maintenance of genetic variation. Evolution (N Y) 68:3166–3183. John Wiley & Sons, Ltd.

Puentes, A., G. Granath, and J. Ågren. 2016. Similarity in G matrix structure among natural populations of Arabidopsis lyrata. Evolution (N Y) 70:2370–2386.

Salinas, S., S. E. Irvine, C. L. Schertzing, S. Q. Golden, and S. B. Munch. 2019. Trait variation in extreme thermal environments under constant and fluctuating temperatures. Philosophical Transactions of the Royal Society B: Biological Sciences 374:20180177.

Schlichting, C. D. 1986. The evolution of phenotypic plasticity in plants. Annu Rev Ecol Syst 17:667–693.

Sgrò, C. M., and A. A. Hoffmann. 2004. Genetic correlations, tradeoffs and environmental variation. Heredity (Edinb) 93:241–248.

Shaw, R. G. 2019. From the past to the future: Considering the value and limits of evolutionary prediction. Am Nat 193:1–10.

Shaw, R. G., and J. R. Etterson. 2012. Rapid climate change and the rate of adaptation: Insight from experimental quantitative genetics. New Phytologist 195:752–765.

Shaw, R. G., and F. H. Shaw. 2014. Quantitative genetic study of the adaptive process. Heredity (Edinb) 112:13–20. Nature Publishing Group.

Sheth, S. N., and A. L. Angert. 2016. Artificial selection reveals high genetic variation in phenology at the trailing edge of a species range. American Naturalist 187:182–193.

Sheth, S. N., M. W. Kulbaba, R. E. Pain, and R. G. Shaw. 2018. Expression of additive genetic variance for fitness in a population of partridge pea in two field sites. Evolution (N Y) 72:2537–2545.

Springate, D. A., and P. X. Kover. 2014. Plant responses to elevated temperatures: A field study on phenological sensitivity and fitness responses to simulated climate warming. Glob Chang Biol 20:456–465.

Stearns, S. C. 1992. The Evolution of Life Histories. Oxford University Press, Oxford.

Stearns, S. C. 1989. Trade-Offs in Life-History Evolution. Funct Ecol 3:259.

Therneau, T. M., and P. M. Grambsch. 2000. Modeling Survival Data: Extending the Cox Mode. Springer US, New York, N.Y.

Torres-Martínez, L., N. McCarten, and N. C. Emery. 2019. The adaptive potential of plant populations in response to extreme climate events. Ecol Lett 22:866–874.

Via, S., and R. Lande. 1987. Evolution of genetic variability in a spatially heterogeneous environment: Effects of genotype–environment interaction. Genet Res 49:147–156. University of Toronto.

Wadgymar, S. M., M. N. Cumming, and A. E. Weis. 2015. The success of assisted colonization and assisted gene flow depends on phenology. Glob Chang Biol 21.

Walsh, B., and M. W. Blows. 2009. Abundant genetic variation + strong selection = multivariate genetic constraints: A geometric view of adaptation. Annu Rev Ecol Evol Syst 40:41–59. Annual Reviews.

Warwick, S., A. Francis, and J. La Fleche. 2000. Guide to wild germplasm of Brassica and allied crops (tribe Brassiceae, Brassicaceae).

Weis, A. E., and W. L. Gorman. 1990. Measuring selection on reaction norms: An exploration of the Eurosta-Solidago system. Evolution (N Y) 44:820–831. John Wiley & Sons, Ltd.

Whitlock, M. C., P. C. Phillips, and K. Fowler. 2002. Persistence of changes in the genetic covariance matrix after a bottleneck. Evolution (N Y) 56:1968–1975.

Willis, J. H., and H. A. Orr. 1993. Increased heritable variation following population bottlenecks: The role of dominance. Evolution (N Y) 47:949.

Wilson, A. J. 2008. Why h2 does not always equal VA/VP? J Evol Biol 21:647–650.

Wilson, A. J., D. Réale, M. N. Clements, M. M. Morrissey, E. Postma, C. A. Walling, L. E. B. Kruuk, and D. H. Nussey. 2010. An ecologist’s guide to the animal model. Journal of Animal Ecology 79:13–26.

Winn, A. A. 2004. Natural selection, evolvability and bias due to environmental covariance in the field in an annual plant. J Evol Biol 17:1073–1083.

Wolak, M. E. 2012. nadiv : an R package to create relatedness matrices for estimating non-additive genetic variances in animal models. Methods Ecol Evol 3:792–796.

Wolak, M. E., and L. F. Keller. 2014. Dominance genetic variance and inbreeding in natural populations. Pp. 104–127 *in* Quantitative Genetics in the Wild. Oxford University Press.

Wood, C. W., and E. D. Brodie. 2015. Environmental effects on the structure of the G-matrix. Evolution (N Y) 69:2927–2940. John Wiley & Sons, Ltd.

Wright, S. 1934. Physiological and evolutionary theories of dominance. Am Nat 68:24–53. Science Press.

Zan, Y., and Ö. Carlborg. 2020. Dynamic genetic architecture of yeast response to environmental perturbation shed light on origin of cryptic genetic variation. PLoS Genet 16:e1008801. Public Library of Science.

