## supporting information for "The capacity for adaptation to climate warming in an annual plant (*Brassica rapa*)"

**Short running title:** Adaptative capacity to climate warming

**Keywords:** adaptation, climate warming, additive genetic variance, dominance genetic variance, plasticity, quantitative genetics, *Brassica rapa*

**CONTENTS**

**Supplementary Methods** 2

*Breeding design—* 2

*Seed number count—* 3

*Phenology models—* 3

*Model specifications for genetic variance models—* 4

*Cross-environment genetic correlations—* 7

**Supplementary Results** 9

*Germination and flowering phenology—* 9

*Relationship between silique number and seed number—* 9

**Supplementary Discussion** 9

*The influence of maternal effects on survivorship—* 9

**Supplementary Literature Cited** 11

**Supplementary Figures** 12

**Supplementary Tables** 20

#### Supplementary Methods

***Breeding design***—

The family-structured experimental population was derived through two generations of greenhouse crosses in 2014/2015 and 2017/2018 (**Fig. S1**). Breeding began with a $F_1$ greenhouse generation of 135 plants, each produced from seed drawn from a unique field parent and crossed in a nested paternal half-sibling design. 45 $F_1$ plants were randomly designated as sires, and each of these were crossed to two unique, randomly chosen dames. This produced an $F_2$ generation of 90 full-sib families, with two families (arbitrarily labeled $M_1$ and $M_2$) nested within each $F_1$ sire. Reciprocal crosses among $F_2$ generation plants produced a set of 62 full-sibling families, such that each parent was mother to half the siblings, and father to the other half (**Fig. S1**). The parents drawn from the $F_2$ generation were chosen to assure the $F_3$ sibships were related as first cousins (two $F_1$ grandparents in common) and/or half-first cousins (one $F_1$ grandparent in common) to several other sibships.

Typically, quantitative genetic studies that estimate additive genetic variance have at least 50 families such that errors arising from a lack of statistical power are reduced (Conner and Hartl 2004). Under a controlled greenhouse environment, we produced 62 different families consisting of reciprocal full-siblings, half-siblings, first cousins, and aunts/uncles (**Fig. S1**). 22 of the 62 families were generated in Summer 2017 under the first maternal group ($M_1$) by Sydney Rotman, while the additional 40 were generated in Summer 2018 under the second maternal group ($M_2$) by Cameron So. A third crossing procedure was completed in Winter 2019 for the third maternal group ($M_3$), but the progeny was not included in this study. The cross using the maternal group $M_1$ only used one seed per sire (**Fig. S1**), thereby excluding $1^{st}$ cousin relationships within the $M_1$ group. The cross using the second maternal group $M_2$ however, used two seeds per sire, creating $1^{st}$ cousin relationships and thus the potential to estimate dominance genetic variance ($V_D$). Alongside $V_D$, this reciprocal full sibship design allowed for isolation of maternal effects. Offspring from each full sibship family carry genotypes drawn from the same two parental genomes, therefore effects attributed by maternal influences could be estimated. Since greenhouse conditions were similar for all $F_2$ dames, estimated maternal effects should be largely due to mitochondrial DNA than the maternal environment.

To generate the $F_3$ experimental generation, seeds from each $F_2$ paternal family of a single maternal group (e.g $M_{1,1}S_1$, $M_{1,2}S_2$ etc) were grown in tall 8x8x20cm containers containing a sandy soil mixture of equal parts HP Mycorrhizae Soil Medium (Pro-Mix ®) and sand. From each $F_2$ paternal family, two seeds were placed within each pot to increase chances of germination. If both seeds germinated, one seedling was randomly removed such that only one plant would remain. After most of the seedlings matured, each of the two plants per $F_2$ paternal family were randomly paired with another plant among the $F_2$ generation. Heterogeneous germination and growth rates led to unequal numbers of pairs within each maternal family, where $M_1$ had 22 families and $M_2$ had 40 families (note: only one plant per paternal family was used in the $M_1$ family). All plants were grown at the University of Toronto's Growth Facilities located at the Earth Sciences Centre.

Each crossing pair was placed within a tray encased by plastic sheeting, allowing water to be filled within the make-shift tub such that the soil columns were well saturated. Plants were grown under greenhouse conditions of 40% relative humidity, 20-26°C day temperatures, 18-24°C night temperatures, and a 15-hour photoperiod to optimize the yield from each plant. Reciprocal crosses were performed by plucking dehiscent anthers using forceps and consequently rubbing the pollen against the stigma of the pollen recipient within each pair. Since *Brassica rapa* is an obligately outcrossing species, it was not necessary to emasculate dames to prevent self-fertilization. Continuously bottom watering the plants and providing water-soluble fertilizer (Nutrite® 20-20-20) twice weekly maintained the flowering period in the parental generation for approximately 2 months, allowing seed production to be optimized. After pollinations were performed, seeds were collected after maturity and stored in coin envelopes at laboratory conditions of initially 21°C, then at 4°C as suggested by (Franks et al. 2019). To minimize the possibility of unintentional pollination events between unpaired individuals by pests, we treated the greenhouse with biological agents first, followed by pesticides (Safer's® End All® and Bayer INTERCEPT™ 60WP). Plants were also spaced well around the greenhouse with branches tied together to prevent further physical contact between inflorescences.

***Seed number count*—**

We counted the number of seeds produced per plant in a subsample of the experimental population to elucidate the relationship between seed pod number and seeds produced. We used the Elmor C3 High Sensitive Seed Counter to expedite this procedure, as it would not be feasible to count hundreds to thousands of seeds without minimizing measurement error. The subset of plants sampled include individuals collected from plots 1, 2, 10, and 11 where half of the plots are split between the heated and ambient environments.

***Phenology models*—**

In the special case of germination time, flowering time, and time to last flower deployment, we used *Cox* mixed-effect regression models (Bradburn et al. 2003) from the R package *survival* version 3.1-8 (Therneau and Grambsch 2000) as phenological data was captured on weekly basis by census and did not accurately capture phenological dates. In these models, an interaction between environment and sibship were set as fixed effects while plot nested under environment was set as a random effect. Phenological dates for each plant were registered as the last Julian calendar day within each census week in **Figure S5** only. Only germinated plants were evaluated in flowering phenology assessments.

***Model specifications for genetic variance models*—**

*Univariate analyses*

We estimated components of genetic variance for each trait per environment/environment. In all cases, we used parameter expanded priors to reduce chain mixing times and to prevent the MCMC chains from being stuck at zero. For Poisson and Gaussian distributed traits, we used a "Fisher" parameter expanded prior (see below). In the special case of lifetime fitness, we attempted a model with a zero-inflated Poisson family specification (Hadfield 2019). Since the output of this model was split into the zero-inflation process and Poisson process, analogous to survival and fecundity, we chose not to include these model outputs. For binary traits, we used priors that followed a $\chi^2$ distribution ($df = 1$) and set the family distribution to "threshold" which uses a probit link (Villemereuil et al. 2012). Note that for binary distributed traits, the residual variance had to be fixed at 1 to estimate other components of variance.

Some of the univariate trait models were conducted using a subset of the total dataset. Fitness components and growth performance traits were subset to the either the proportion of plants that germinated or germinated and flowered. **Figure S4** is an expanded graphical model that summarizes the life cycle based on the traits collected and how fitness components or traits potentially related to fitness were subset from the dataset. For example, survival included all the plants in an environment and was divided binarily as either successful survival to flowering or failure. On the other hand, fecundity only included plants that produced at least 1 inflorescence and was represented by the number of siliques produced. This procedure was used to prevent trait distributions from being over-inflated at zero and to prevent violations of model assumptions.

To ensure each model cleared all diagnostic criteria (i.e. adequate convergence, >1000 effective sample sizes, autocorrelation <0.1), we ran binary trait models for 1,100,000 iterations with a 100,000 burn-in storing samples every 500 iterations. Non-binary trait models were run for 2,100,000 iterations with a 100,000 burn-in storing samples every 1000 iterations. In the special case of trait inflorescences, we ran the model for 8,500,000 iterations with a 500,000 burn-in storing samples every 4000 iterations. This was necessary to reduce autocorrelation between posterior estimates. Since stored posteriors produced bimodal distributions, likely as a result of priors pulling estimates toward zero, we obtained estimates of variance using posterior means rather than posterior modes. We used multiple priors (see below) in attempt to produce unimodal posteriors, but all posterior distributions tailed towards zero.

For each environment, we obtained estimates of additive genetic variance ($V_A$), dominance genetic variance ($V_D$), maternal effect variance ($V_m$) and residual variance ($V_R$) using posterior means. The rate of adaptation ($\Delta_E \overline{W}$) and narrow sense heritability ($h^2$) (**Equation 2&3**) were calculated from these estimates on both the latent and data scale. We opted not to use deviance information criterion (DIC) values to evaluate the significance of dominance variance, as comparing DIC values is not a perfect model selection criterion especially for non-Gaussian models (Hadfield 2019). After genetic variance parameters were estimated, test of significance between the two environments were assessed by using 95% confidence intervals. Confidence intervals that did not overlap indicated significance.

$$\Delta_E \bar{W} = \frac{V_A(W)}{\bar{W}} \quad [2]$$

$$h^2 = \frac{V_A}{V_A + V_D + V_m + V_R} \quad [3]$$

*Prior for Poisson/Gaussian Traits:*

G1: V = 1, nu = 1, alpha.mu = 0, alpha.V = 1000

G2: V = 1, nu = 1, alpha.mu = 0, alpha.V = 1000

G3: V = 1, nu = 1, alpha.mu = 0, alpha.V = 1000

R: V = 1, nu = 0.002

*Alternative Priors for Poisson/Gaussian Traits:*

G1: V =1, nu = 0.002, alpha.mu = 0, alpha.V = 1000

G2: V =1, nu = 0.002, alpha.mu = 0, alpha.V = 1000

G3: V = 1, nu = 0.002, alpha.mu = 0, alpha.V = 1000

R: V = 1, nu = 0.002

*Priors for Binary Traits:*

G1: V = 1, nu = 1000, alpha.mu = 0, alpha.V = 1

G2: V = 1, nu = 1000, alpha.mu = 0, alpha.V = 1

G3: V = 1, nu = 1000, alpha.mu = 0, alpha.V = 1

R: V =1, fix = 1

*Multivariate analyses*

We attempted to construct a **G** matrix between survival and fecundity to estimate the genetic correlation between fitness components. The bivariate model between survival and fecundity included all individuals in each environment. We attempted a model with a prior that weakly centred the genetic correlation around zero (see below). Models were run for 2,100,000 iterations with a 100,000 burn-in storing samples every 1000 iterations and subsequently checked for any convergence issues, high autocorrelation, or low effective sample size. In all cases, models including dominance variance were not able to pass diagnostic tests. Therefore, we estimated the genetic correlation between fitness components using an approach of resampling posteriors (Grieshop et al. 2021; see section below for details).

*Prior 1 for G-matrix model:*

G1: V = diag(2)*0.02, nu = 3, alpha.mu = c(0,0), alpha.V = diag(c(1000,1000))

G2: V = diag(2)*0.02, nu = 3, alpha.mu = c(0,0), alpha.V = diag(c(1000,1000))

G3: V = diag(2)*0.02, nu = 3, alpha.mu = c(0,0), alpha.V = diag(c(1000,1000))

R: V = diag(2)*0.02, nu = 3, fix = 2

*Prior 2 for G-matrix model:*

G1: V = diag(2), nu = 2, alpha.mu = c(0,0), alpha.V = diag(c(1,1))

G1: V = diag(2), nu = 2, alpha.mu = c(0,0), alpha.V = diag(c(1,1))

G1: V = diag(2), nu = 2, alpha.mu = c(0,0), alpha.V = diag(c(1,1))

R: V = diag(2), nu = 2, fix = 2,

***Cross-environment genetic correlations***—

We estimated the genetic correlation for trait expression across the ambient and heated environments by resampling breeding values from each within-environment model. Specifically, we resampled $F_2$ (parents of the $F_3$ experimental population) breeding values from 2000 stored posteriors and subsequently generated a distribution of cross-environment genetic correlation values using **Equation 4**, where *i* and *j* represent either the ambient or heated environment. This approach incorporated the uncertainty around the breeding values into the estimation of *r* and its credibility intervals, thus permitting tests of significance in a conservative manner despite additive covariances not being accounted for in the model. (Grieshop et al. 2021).

$$r_{Aij} = \frac{Cov_{Aij}}{\sqrt{(V_{Ai})(V_{Aj})}} \qquad [4]$$

We estimated the cross-environment genetic correlation ($r_A$) using the approach above because models that included data from both environments and that attempted to estimate the 3 components of cross-environment covariance had issues of convergence and autocorrelation. In reference, these ‘full’ models specified a “us(environment):animal” structure in the model, where “us()” estimates the covariance between environments (Hadfield 2019). To provide confidence that our approach of estimating $r_A$ from separate environmental models did not differ from the approach using cross-environment models, we estimated $r_A$ using both approaches but in models that excluded the dominance variance component. By removing dominance variance and only estimating within-environment variances using the “idh()” function, we were able to produce models passing diagnostic tests. In these cross-environment models, we also set environment and plot nested within environment as a fixed effect. Models were run for 2,100,000 iterations with a 100,000 burn-in storing samples every 1000 iterations.

*Parallel processes and stacking MCMC chains*

For models of longer estimated run-times, we ran parallel models and stacked MCMC chains using a function provided by Maurizio et al. (2018). Based on the number of cores available on the server, we were able to reduce run-times by a factor of cores available.

```
mcmc.stack <- function (coda.object, ...){
 ## This function is from Will; also part of BayesDiallel
 if (inherits(coda.object, "mcmc")) {
  return(coda.object)
 }
 if (!inherits(coda.object, "mcmc.list")) {
  stop("Non-mcmc object passed to function\n")
 }
 chain <- coda.object[[1]]
 for (i in 2:nchain(coda.object)) {
  chain <- rbind(chain, coda.object[[i]])
 }
 as.mcmc(chain)
}
```

### Supplementary Results

##### *Germination and flowering phenology*—

Warming advanced the timing of germination ($\chi^2$ = 190.90; $p$ < 2.2e-16; β = exp(0.38)) relative to the ambient environment. The median germination time differed by 10 days between the two environments (**Fig. S5A**). This resulted in plants deploying their first flower earlier in the heated environment ($\chi^2$ = 431.87; $p$ < 2.2e-16; β = exp(7.80)), and consequentially completing their flowering season sooner ($\chi^2$ = 380.08; $p$ < 2.2e-16; β = exp(1.07)). Median time to first flower differed by 5 days between environments, and 7 days for time to flowering completion (**Figs. S5B&C**). Overall, plant exposure to warmer climatic conditions imposed by open-top greenhouse chambers and artificial warming ultimately shifted the timing of germination and flowering phenology to earlier dates compared to ambient conditions**.**

##### *Relationship between silique number and seed number*—

We used Pearson's correlation $r$ to evaluate the relationship between silique production and the number of seeds produced per plant. We found a strong relationship between the two fitness-related traits ($R^2$ = 0.831; **Fig. S8**), thus reassuring our choice of silique number as a good proxy for absolute fitness where otherwise the number of seeds produced would represent the seed to seed development of progeny.

### Supplementary Discussion

##### *The influence of maternal effects on survivorship*—

Maternal effects in plants have largely been known to affect seed size, dormancy dynamics, early seedling survival (Singh et al. 2017), and if present, can obscure genetic variation and slow adaptative evolution (Roach and Wulff 1987). The contribution of maternal effects to survival and survival decomposed into two separate components, overwintering survival and spring-summer survival, was negligible in both ambient and heated conditions. Low maternal effects contributing to survival variance may indicate maternal identity is trivial to adaptation to future climate warming and has limited contribution to lifetime fitness.

However, failure to detect larger maternal effects may also be a result of conducting crosses in a controlled greenhouse environment to produce the experimental generation (Wadgymar et al. 2018a). By providing ample water resources, supple nutrients, and a stable climatic condition to the generations prior to the experimental population, the heterogeneity of finding such conditions in the wild is not properly represented (Poorter et al. 2016). As such, the stable maternal environment likely reduced the variance of endosperm resources provided to offspring among pedigreed families. If the greenhouse maternal environment matched field conditions, then transgenerational plasticity may have strengthened maternal effects contributing to phenotypic variance (Mousseau and Fox 1998; Galloway 2005). Predictable offspring environments can allow maternal plants to adjust their investment into seed resources and other traits to best maximize offspring fitness (Herman et al. 2012). By chance, seeds can disperse into habitats that misalign with their maternal plant and result in reduced fitness. The degree to which transgenerational plasticity is expressed has consequences on fitness and fitness variation.

Assuming the parental generation was grown and crossed in the field without error, stronger maternal effects in survival and subsequent life-history traits may be observed. For example, a reciprocal transplant study across an elevation gradient revealed transgenerational plasticity influences fitness-related traits including seed and seedling performance with clinal patterns (Wadgymar et al. 2018a). However, maternal effects through transgenerational plasticity can be insignificant to legacy effects left by extreme environmental conditions such as drought (De Long et al. 2019). It is important to note that maternal effects deriving from transgenerational plasticity is a special form of phenotypic plasticity (Galloway and Etterson 2007), and if climate change decouples the ability to predict offspring environments, it can be maladaptive (Visser 2008). Regardless, even if fitness is improved by adaptive transgenerational plasticity, long-term population persistence is still dependent on adaptive evolution rather than simple changes to mean fitness (Shaw and Shaw 2014).Therefore, assessing the underlying additive genetic variance of transgenerational plasticity will further our understanding of adaptation to future climate warming and predicting population persistence (Kuijper and Hoyle 2015; Donelson et al. 2018).

#### Supplementary Literature Cited


Bradburn, M. J., T. G. Clark, S. B. Love, and D. G. Altman. 2003. Survival Analysis Part II: Multivariate data analysis – an introduction to concepts and methods. British Journal of Cancer 89:431–436.

Conner, J. K., and D. L. Hartl. 2004. A Primer of Ecological Genetics. Sinauer Associates, Inc., Sunderland, USA.

Franks, S. J., M. R. Sekor, S. Davey, and A. E. Weis. 2019. Artificial seed aging reveals the invisible fraction: Implications for evolution experiments using the resurrection approach. Evolutionary Ecology 33:811–824.

Grieshop, K., P. L. Maurizio, G. Arnqvist, and D. Berger. 2021. Selection in males purges the mutation load on female fitness. Evolution Letters 5:328–343. John Wiley & Sons, Ltd.

Hadfield, J. 2019. MCMCglmm Course Notes.

Maurizio, P. L., M. T. Ferris, G. R. Keele, D. R. Miller, G. D. Shaw, A. C. Whitmore, A. West, C. R. Morrison, K. E. Noll, K. S. Plante, A. S. Cockrell, D. W. Threadgill, F. Pardo-Manuel de Villena, R. S. Baric, M. T. Heise, and W. Valdar. 2018. Bayesian Diallel Analysis Reveals Mx1-Dependent and Mx1-Independent Effects on Response to Influenza A Virus in Mice. G3 Genes|Genomes|Genetics 8:427–445. Oxford Academic.

Therneau, T. M., and P. M. Grambsch. 2000. Modeling Survival Data: Extending the Cox Mode. Springer US, New York, N.Y.

Villemereuil, P. De, J. A. Wells, R. D. Edwards, and S. P. Blomberg. 2012. Bayesian models for comparative analysis integrating phylogenetic uncertainty. BMC Evolutionary Biology 12:102.

#### Supplementary Figures

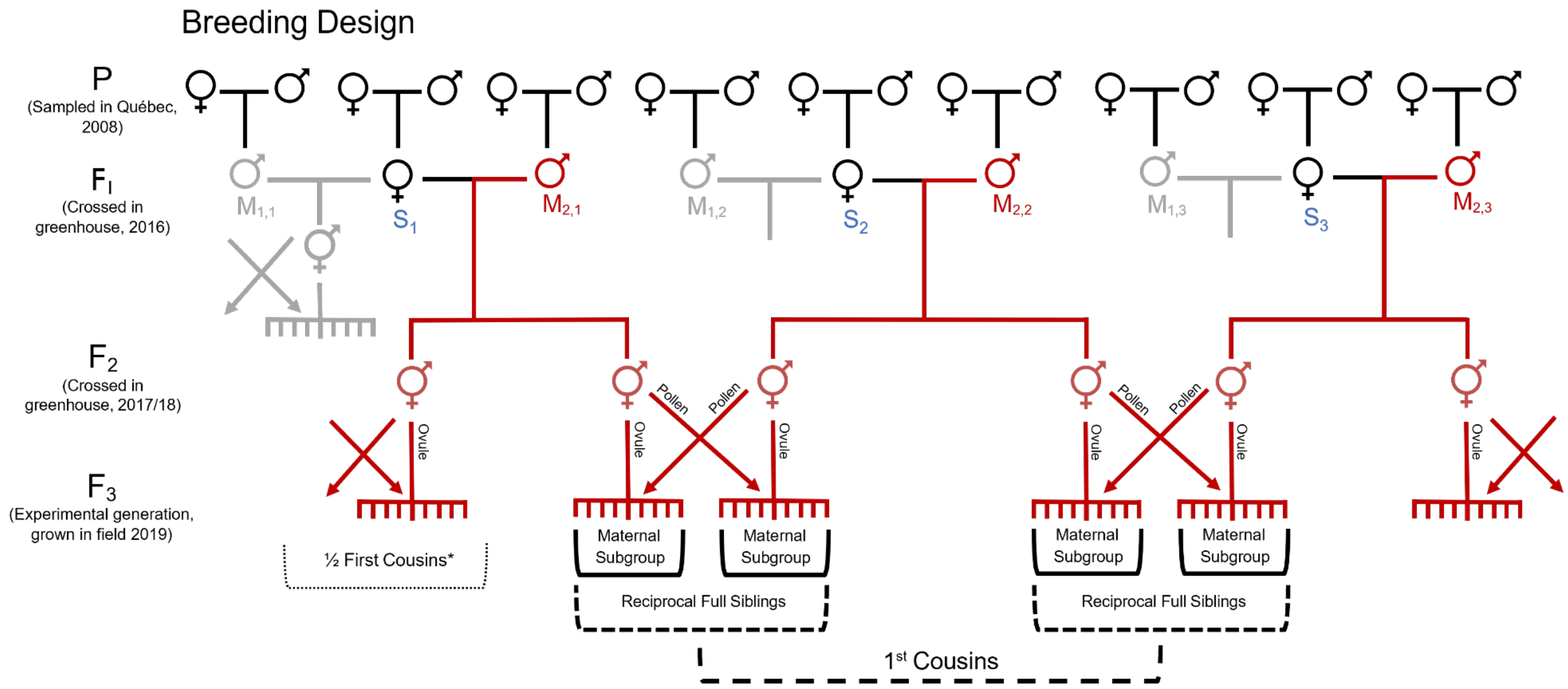


**Figure S1:** A pedigree showing how the experimental generation ($F_3$) was produced from the initial population (P) collected from Saint-Georges, Québec in 2008. **This pedigree only displays crosses from the second maternal group ($M_2$) coloured in red**, in which dams ($F_1$ grandparents) were initially crossed with 45 sires ($S_1$ through $S_{45}$). The progeny ($F_2$) of the first grandparental cross were then used to generate the experimental population consisting of reciprocal full siblings, 1$^{st}$ cousins, and half-sibling relationships. Note that the half-siblings are not shown because another maternal group ($M_1$) is not included. *Half 1$^{st}$ cousin relationships are formed from progeny that share the same grandfather (although not shown, the grey progeny was created by reciprocal crosses identical to the $F_2$ crosses, within their respective maternal group (*i.e.,* $M_1$ or $M_2$). Figure borrowed from So et al. (in press).

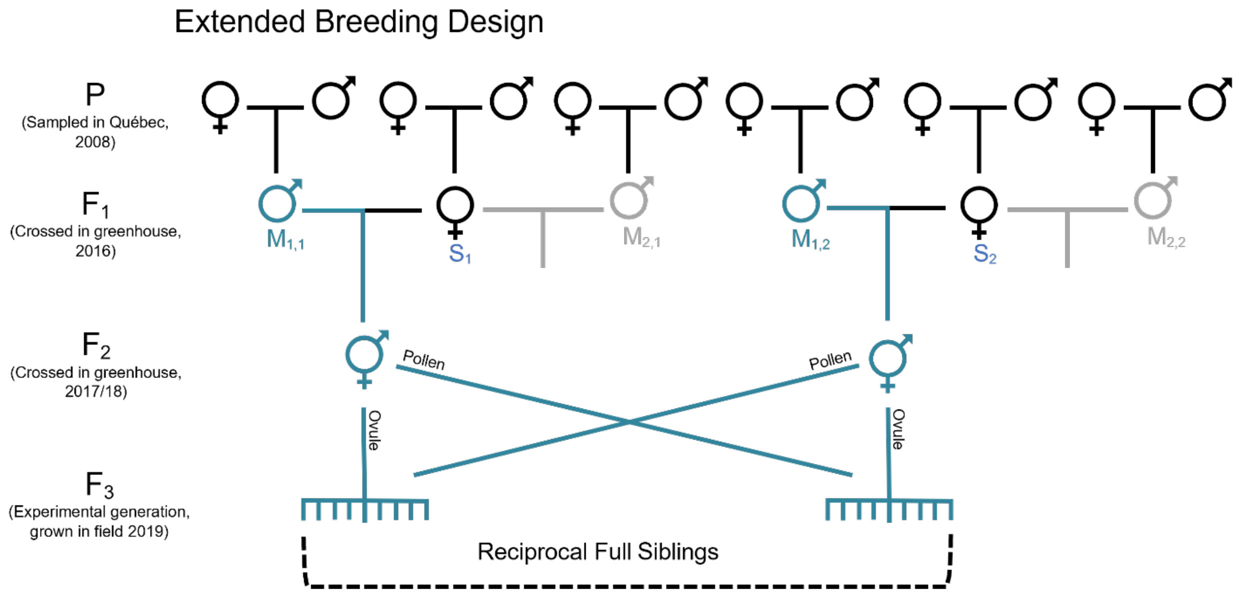


**Figure S2:** A pedigree showing how the experimental generation ($F_3$) was produced and derived from the initial population (P) collected from Saint-Georges, Québec in 2008. **This pedigree shows the crosses within the first maternal group ($M_1$) coloured in turquoise**. Unlike the second maternal group, this maternal group does not contain 1st cousin relationships for estimating the genetic variance component of dominance. Figure borrowed from So et al. (in press).

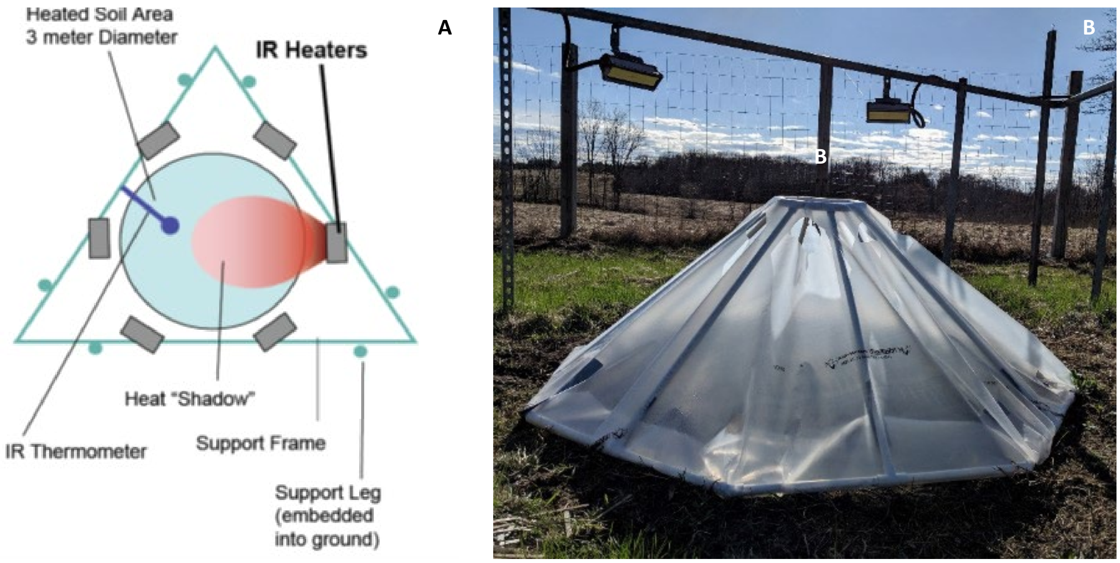


**Figure S3: (A)** Six heaters emit infrared to increase the ambient temperature by a set number of degrees. The triangular frame supports six infrared heaters such that a 3-metre diameter plot is heated. **(B)** An open-top chamber deployed on one of six heated plots in late winter-early spring to impose an early spring seasonality as predicted by the IPCC.

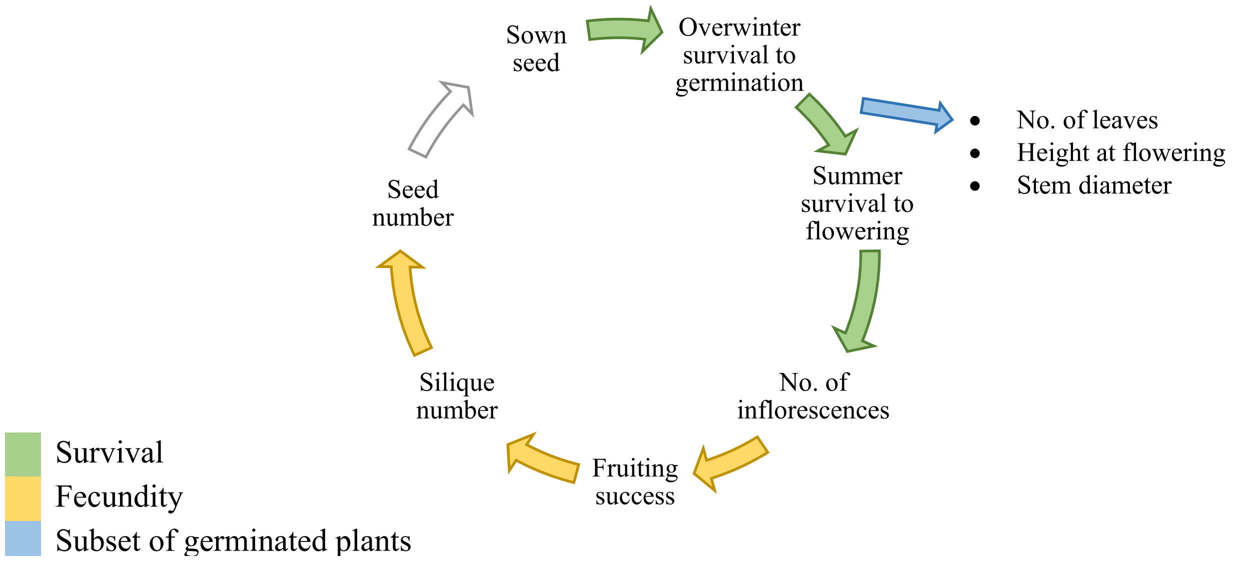


**Figure S4:** The life cycle of *Brassica rapa* constructed using collected fitness component traits. The green arrows represent the fitness component overwintering survival to flower production, while the yellow arrows represent the fitness component of fecundity of successfully flowering plants.

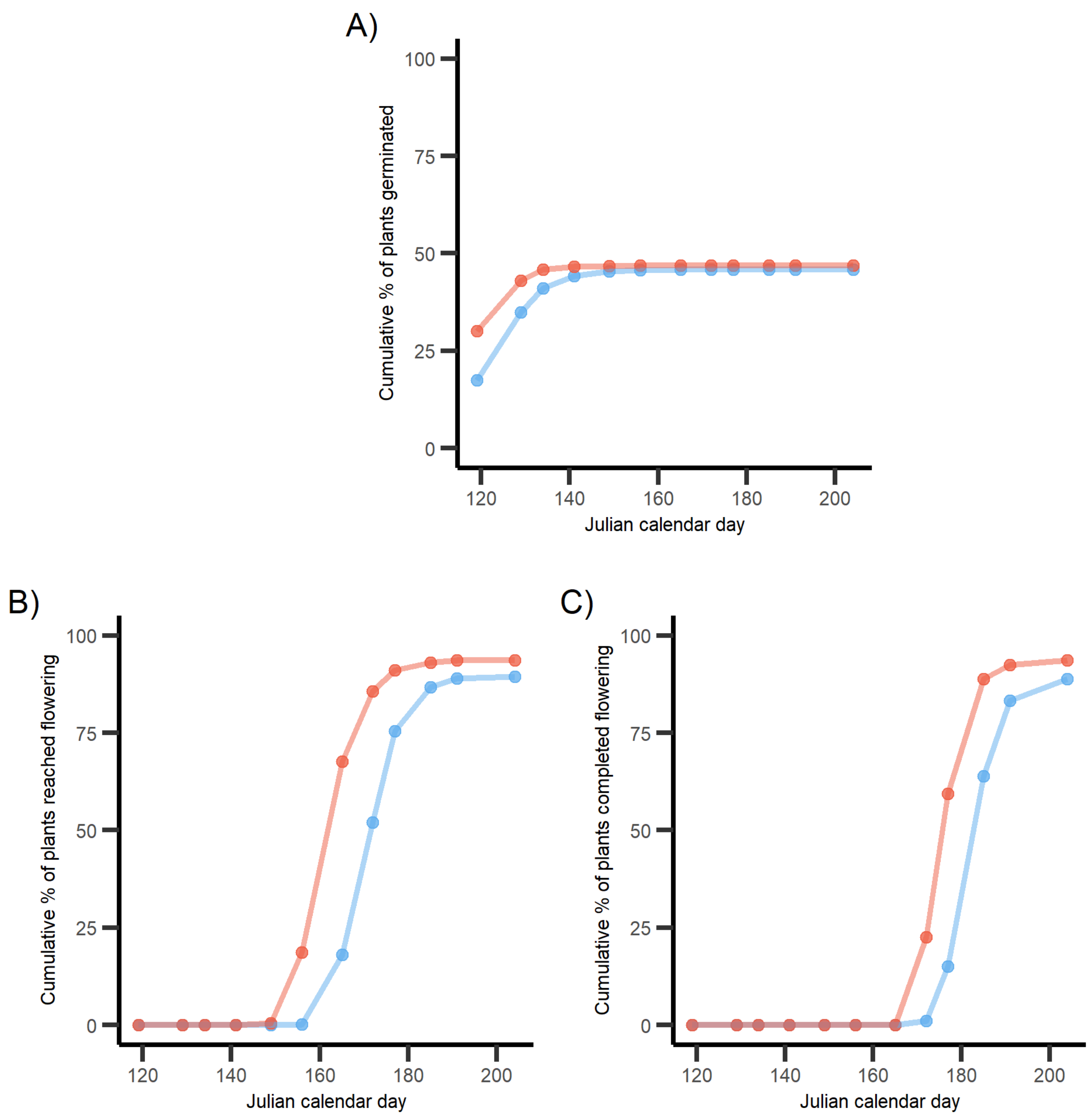


**Figure S5:** Cumulative proportion of plants that **(A)** germinated, **(B)** germinated and reached the flowering stage, and **(C)** completed flowering, plotted against the last Julian calendar day within a census. The ambient environment is represented by the blue colouration, while the heated environment is represented by red. Exposing *Brassica rapa* to a future climate scenario influenced plants to flower earlier than the ambient environment.

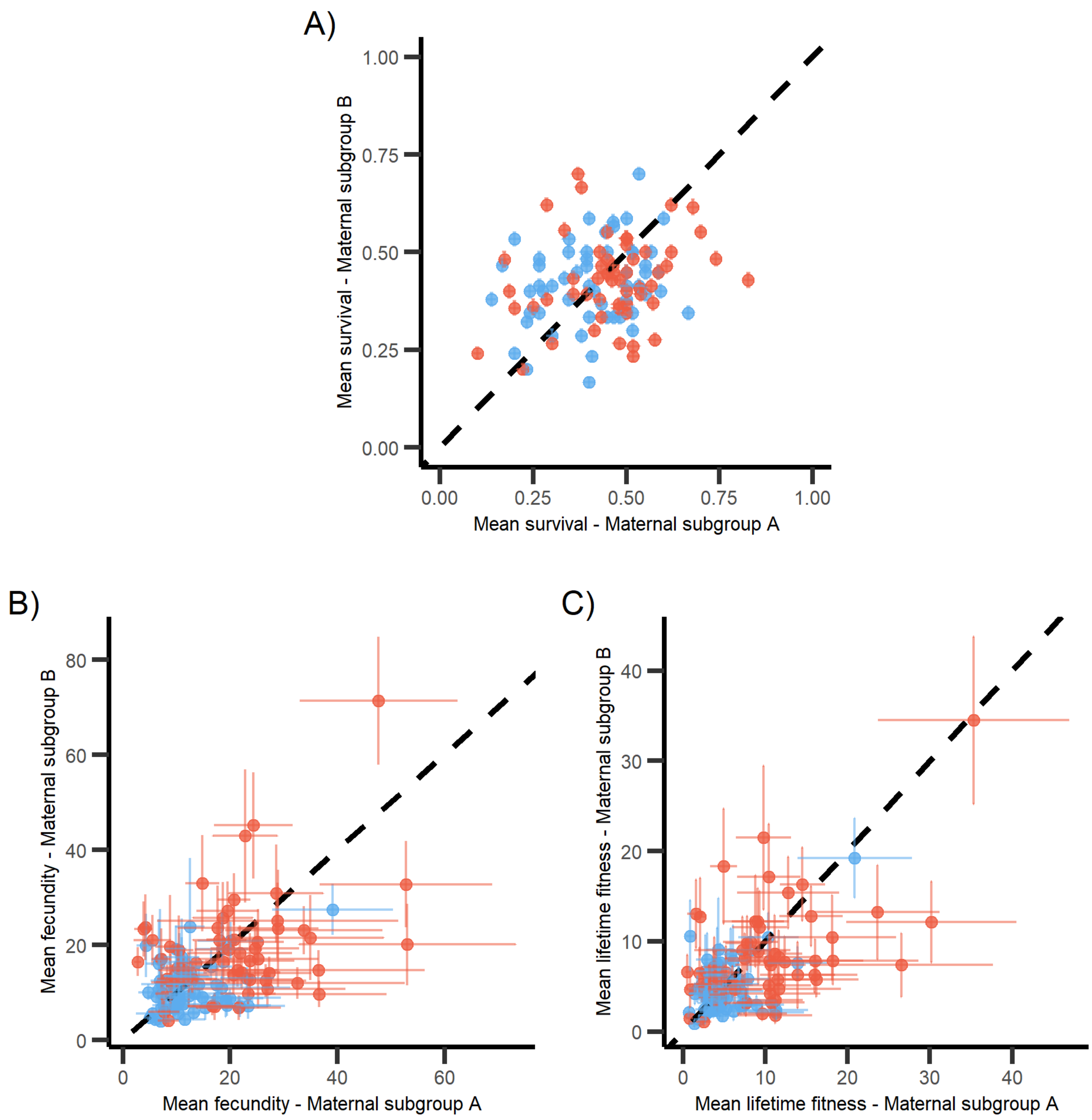


**Figure S6:** Within-family maternal variance displayed by plotting mean trait values of a maternal subgroup against the other maternal subgroups. The heated environment is represented by red dots while the ambient environment is shown in blue. Error bars are the standard error of the mean. Deviation from the 1:1 line indicates maternal effects contributing to the variance in **(A)** overwintering survival to flowering, **(B)** fecundity of flowering plants, and **(C)** lifetime fitness.

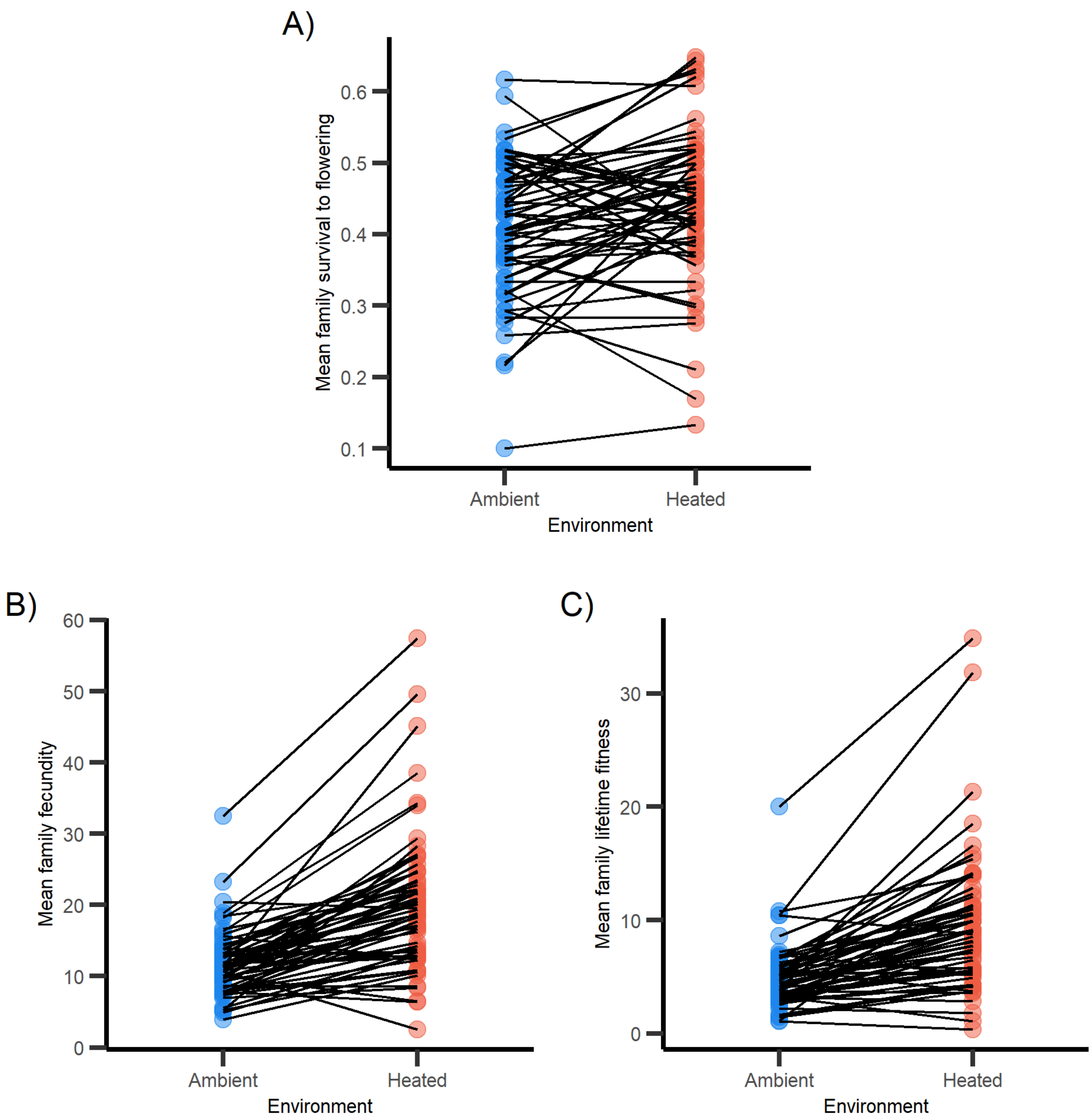


**Figure S7**: Reaction norms of sibship means between the two environments for **(A)** overwintering survival to flowering, **(B)** fecundity of flowering plants, and **(C)** lifetime fitness.

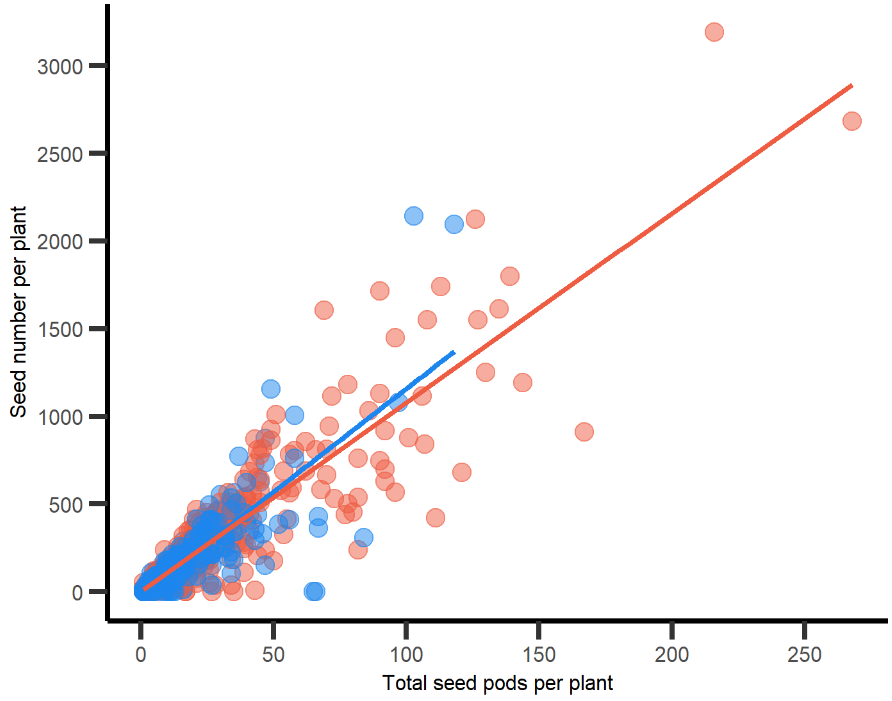


**Figure S8:** Evaluation of silique (seed pod) number as a proxy for absolute fitness measured by seed number (total $R^2 = 0.831$)

### Supplementary Tables

**Table S1**: A summary of trait means and probabilities per environment group, followed with significance testing of the environment using Type II ANOVAs applied for reproductive components. Standard errors are provided following trait means and probabilities.

| Trait (*i*) | Filter | GLMM Family | Ambient $\mu$ or $p$ (SE) | Heated $\mu$ or $p$ (SE) | Type II Wald $\chi^2$ | Environment (*P*-value) |
|---|---|---|---|---|---|---|
| ***Reproductive components*** | | | | | | |
| Overwintering survival | Germinated | Gaussian | 0.46 (0.038) | 0.47 (0.037) | $0.063_{(1,6802)}$ | 0.80 |
| Spring-summer survival | Germinated | Gaussian | 0.90 (0.011) | 0.94 (0.0072) | $12.76_{(1,3161)}$ | **3.53e-04** |
| Fruiting success | Germinated | Gaussian | 0.993 (0.0023) | 0.996 (0.0016) | $1.15_{(1,2902)}$ | 0.28 |

**Table S2: Survival and reproductive component** estimations of additive genetic variance ($V_A$), dominance genetic variance ($V_D$), maternal effects ($V_m$), narrow-sense heritability ($h^2$), and adaptive potentials ($\Delta_E \bar{z}$). The variances of Bernoulli traits are shown on both the latent and data scale, while the 95% confidence intervals are only provided on the data scale.

| | | $V_A$ | | $V_D$ | | $V_M$ | | $h^2$ | | $\Delta_E \bar{Z}$ | |
|---|---|---|---|---|---|---|---|---|---|---|---|
| | | Ambient | Heated | Ambient | Heated | Ambient | Heated | Ambient | Heated | Ambient | Heated |
| Overwintering survival | Latent | 0.0219 | 0.0301 | 0.0496 | 0.117 | 0.0269 | 0.0351 | 0.0198 | 0.0256 | - | - |
| | Data | 1.63e-3 | 2.09e-3 | 3.62e-3 | 7.95e-3 | 2.01e-3 | 2.45e-3 | 6.55e-3 | 8.40e-3 | 0.0139 | 0.0176 |
| | CI-Lo | 1.61e-10 | 9.10e-11 | 3.03e-8 | 2.12e-9 | 2.66e-10 | 2.08e-9 | 6.46e-10 | 3.64e-10 | 1.37e-9 | 7.76e-10 |
| | CI-Hi | 5.25e-3 | 7.05e-3 | 0.116 | 0.0202 | 4.06e-3 | 4.84e-3 | 0.0210 | 0.0282 | 0.0447 | 0.0601 |
| Spring-summer survival | Latent | 0.0536 | 0.102 | 0.101 | 0.269 | 8.95e-3 | 0.110 | 0.451 | 0.0672 | - | - |
| | Data | 1.68e-3 | 1.58e-3 | 3.28e-3 | 4.75e-3 | 2.83e-4 | 1.78e-3 | 0.0113 | 0.0150 | 0.0138 | 0.0170 |
| | CI-Lo | 3.65e-9 | 2.45e-11 | 4.22e-9 | 2.59e-8 | 5.54e-9 | 1.70e-9 | 2.43e-8 | 2.67e-10 | 2.96e-8 | 3.03e-10 |
| | CI-Hi | 5.04e-3 | 5.76e-3 | 0.0121 | 0.0155 | 1.11e-3 | 4.45e-3 | 0.0335 | 0.0540 | 0.0410 | 0.0611 |
| Fruiting success | Latent | 0.0578 | 0.104 | 0.0735 | 0.0780 | 0.0643 | 0.0522 | 0.0457 | 0.0794 | - | - |
| | Data | 1.33e-4 | 1.40e-4 | 1.94e-4 | 1.20e-4 | 1.67e-4 | 7.62e-5 | 4.09e-3 | 5.91e-3 | 4.22e-3 | 6.04e-3 |
| | CI-Lo | 4.52e-11 | 2.55e-12 | 5.04e-11 | 6.76e-12 | 9.27e-10 | 4.79e-12 | 1.57e-9 | 1.28e-10 | 1.62e-9 | 1.31e-10 |
| | CI-Hi | 5.26e-4 | 4.92e-4 | 7.89e-4 | 4.73e-4 | 6.48e-4 | 3.02e-4 | 0.0155 | 0.0193 | 0.0161 | 0.0198 |

**Table S3:** Cross-environment additive genetic correlations ($r_A$) and $p$ values signifying any differences of $r$ from 0 for survival and reproductive components.

| Trait | Genetic Correlation ($r_A$) | Significance ($p$-value) |
|---|---|---|
| Overwintering survival | 0.0307, (-0.163, 0.208) | 0.845 |
| Spring-summer survival | 0.00247, (-0.172, 0.203) | 0.912 |
| Fruiting success | -0.0260, (-0.184, 187) | 0.986 |

**Table S4:** Additive genetic correlations ($r_A$) between survival and fecundity for each of the environments. 95% confidence intervals are provided. The *p*-values indicate significant differences from 0.

| Environment | Genetic Correlation ($r_A$) | Significance (*p*-value) |
|---|---|---|
| Ambient | 0.0247<br>(-0.108, 0.151) | 0.70 |
| Heated | 0.0254,<br>(-0.104, 0.160) | 0.73 |
